# Multiplicity of Type 6 Secretion System toxins limits the evolution of resistance

**DOI:** 10.1101/2024.07.30.605577

**Authors:** William P. J. Smith, Ewan Armstrong-Bond, Katharine Z. Coyte, Christopher G. Knight, Marek Basler, Michael A. Brockhurst

## Abstract

The bacterial Type 6 Secretion System (T6SS) is a toxin-injecting nanoweapon that mediates competition in plant- and animal-associated microbial communities. Bacteria can evolve *de novo* resistance against T6SS attacks, but resistance is far from universal in natural communities, suggesting key features of T6SS weaponry may act to limit its evolution. Here, we combine eco-evolutionary modelling and experimental evolution to examine how toxin type and multiplicity in *Acinetobacter baylyi* attackers shape resistance evolution in susceptible *Escherichia coli* competitors. In both our models and experiments, we find that combinations of multiple distinct toxins limit resistance evolution by creating genetic bottlenecks, driving resistant lineages extinct before they can reach high frequency. We also show that, paradoxically, single-toxin attackers often drive the evolution of cross-resistance, protecting bacteria against unfamiliar toxin combinations, even though such evolutionary pathways were inaccessible against multi-toxin attackers. Our findings indicate that, comparable to antimicrobial and anticancer combination therapies, multi-toxin T6SS arsenals function to limit resistance evolution in competing microbes. This helps us to understand why T6SSs remain widespread and effective weapons in microbial communities, and why many bacteria T6SS-armed encode functionally diverse anti-competitor toxins.

**Significance:** Toxin secretion systems, such as T6SSs, are widely used by bacteria to inhibit competing microorganisms. Here, we show that the secretion of multiple toxins in combination can suppress the evolution of resistance to the T6SS, rationalising its continued widespread use across diverse communities. Our work shows that principles of combination therapy—well known in the context of antimicrobial, antiviral and anticancer therapies—also apply in the context of microbial warfare, helping to explain why many bacteria maintain diverse T6SS toxin arsenals. Resistance suppression is also a technologically useful property of T6SS toxin cocktails, which could be harnessed as part of future biocontrol or biotherapeutic applications, using live T6SS-armed bacteria to limit the growth of problem microorganisms.

## Introduction

The Type 6 Secretion System (T6SS) is a molecular harpoon gun, used extensively by diderm bacteria to deliver effector proteins into diverse target cells, including both diderm and monoderm bacteria^1^, fungi^2^, eukaryote predators^3,4^, and animal hosts^5^. While T6SSs are highly versatile, with roles spanning virulence^6^, host manipulation^7^, and metal scavenging^8,9^, their primary function appears to be as anti-competitor weapon systems^10^. By injecting toxic effector proteins (referred to hereafter as “toxins”) into rivals, T6SSs enable bacteria to eliminate competing strains and species, and so claim space, nutrients and genetic material^11,12^. In this capacity, T6SS killing is a major determinant of bacterial strains’ success: T6SSs play crucial ecological roles in multiple host-associated microbial communities, including within the mammalian gastro-intestinal tract^13^, and in the plant rhizosphere and phytosphere^14^. Additionally, T6SSs are used by human pathogens to eliminate competing commensals, and thereby establish^15^ or consolidate^16^ infections in various body sites.

A distinctive feature of T6SSs is that they are inherently mechanical weapons^17^: by piercing cell membranes and translocating toxins directly into target cells, they avoid reliance on specific surface receptors or membrane transporters^18^. This mechanical mode of action also gives the T6SS the unusual ability to translocate multiple, functionally-different toxins at the same time^19^. The ecological and clinical importance of T6SSs has sparked recent interest in how microbes might protect themselves from these potent, multi-pronged toxin attacks^20–23^. Canonically, protection against T6SS attacks is conferred through the expression of immunity proteins: co-expressed with their cognate toxins, these immunity factors protect T6SS-wielding cells both from their own toxins, and from ‘friendly fire’ from neighbouring kin^19^. Intriguingly, various bacteria retain “orphan” immunity proteins for toxins they themselves do not produce, underscoring their value as a protection against attack by competitors^24–26^.

However, a bacterium’s defence against T6SS toxins is not limited to possession of immunity proteins. In the last 7 years, multiple alternate forms of defence have been reported, demonstrating T6SS protection via biofilm^27^ or microcolony formation^28^, cell envelope stress responses^29^, capsule formation^30,31^, toxin-specific defence pathways^32^, or through loss of susceptibility factors^33^. In some cases, these resistance mechanisms can obviate the need for immunity proteins altogether^29^; recent work has also demonstrated microbes’ potential to evolve spontaneous resistance to T6SS attacks^34^. However, because these studies have generally focused on resistance mechanisms against T6SS attackers with fixed toxin arsenals, the role of specific toxins or toxin combinations in determining weapon susceptibility and evolutionary responses has remained thus far uncharacterised.

Despite the potential for resistance against T6SS to evolve *de novo*, in natural communities T6SSs appear to retain their utility over ecological and evolutionary timescales^25,35^, suggesting that there are key features of T6SSs that may act to limit resistance evolution. In particular, T6SSs’ capacity to inject multiple, functionally different toxins in combination might be a key determinant of resistance evolution potential. First, different T6SS toxins might be intrinsically more or less easy to resist than others, depending on their mode of toxicity. Second: analogous to anticancer^36^, antiviral^37^, and antibiotic combination therapies^38^, multi-toxin secretion might limit resistance compared with mono-toxin secretion, requiring a more complex or more costly combination of resistance mutations for resistance to emerge and spread^39,40^.

Here, we test the hypothesis that T6SS toxin type and multiplicity act to limit resistance evolution. We develop a new agent-based model of multi-toxin T6SS warfare, which predicts that multi-toxin attackers can suppress the evolution of T6SS resistance: both by limiting the range of viable resistance mutations (those subject to positive natural selection), and ii) by driving evolving lineages extinct with greater frequency. Counterintuitively, our models also predict that single-toxin attackers can drive the evolution of both cross-resistance (protection against previously unencountered toxins) and multi-resistance (protection against toxin combinations). We then test these predictions using experimental evolution, passaging *E. coli* bacteria against *Acinetobacter baylyi* attackers armed with single or multiple common T6SS toxin types. Using combinatorial competition assays, we test for evolved resistance against different toxin types: while multi-toxin attackers result in high extinction rates and minimal resistance, single-toxin attackers select frequently for cross-resistance to other toxins. Overall, our experimental findings confirm our model predictions, demonstrating that both T6SS toxin type and multiplicity act to limit the evolution of resistance in susceptible bacterial populations.

## Results

### An agent-based model describes cellular responses to T6SS toxins and toxin combinations

To investigate how toxin multiplicity impacts T6SS resistance evolution, we began with an agent-based computer model of T6SS warfare, based on the open-source software *Cellmodeller*^41^. This framework has been used previously to study T6SS toxin evolution^42^, T6SS regulation^43^, collective protection^27^, and T6SS function alongside other weapons^44^. Crucially, it allows one to study cell-cell interactions in a spatially structured bacterial community featuring realistic cell packing and biomechanical forces. Community spatial structure is centrally important in determining the outcomes of T6SS competition^23,42,45^, making this model a natural starting point for our investigations.

Our model presents a simple yet flexible description of sensitivity to multiple T6SS toxins, secreted alone or in combination (see Figs. 1 and S1, and Materials and Methods for a full description). To the best of our knowledge, ours is the first model of T6SS warfare to consider multiple toxins. Notionally, it can represent different types of effectors with different potencies, doses and mechanisms of toxicity (although such mechanisms are not explicitly modelled). Briefly, our model is as follows: i) we consider populations of bacterial cells placed randomly on a 2-D surface; ii) cells are represented as elongating rods that push on one another as they grow and divide; iii) cells are of one of two basic types: T6SS+ attackers, and T6SS-target cells with variable levels of resistance (Fig. 1A); iv) attacker cells randomly fire T6SS needles at a specific rate, translocating toxins into neighbouring cells (attackers are completely immune to these toxins); v) susceptible cells incur cellular damage proportional to a) their exposure to translocated toxins and b) their individual sensitivities to those toxins, which are controlled by sensitivity parameters ai. For reference, we summarise model variables and parameters, along with their values and sources, in Tables S1 and S2 respectively.

**Fig. 1:**
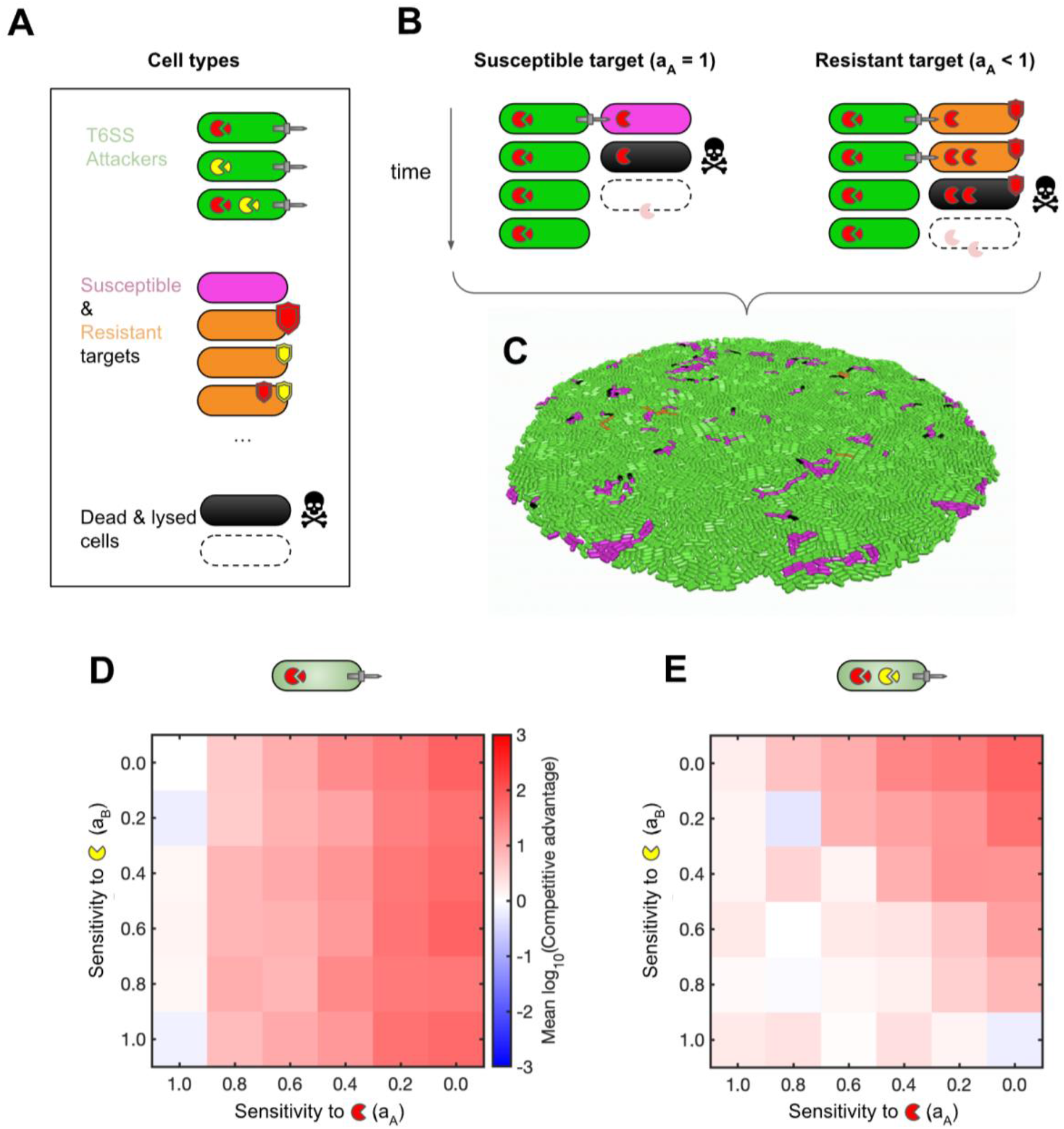
Agent-based modelling of T6SS warfare predicts that multi-toxin attackers limit resistance evolution in susceptible competitors. **(A)** Cell types in this model include T6SS attackers of variable toxin arsenal (top; toxins shown as coloured pac-men), T6SS-susceptible cells with independently variable sensitivity to these toxins (middle; resistance shown as coloured shields). T6SS intoxication ultimately leads to target cell death (bottom). **(B)** Sensitivity to each toxin i is controlled via parameter a_i_; a cell’s sensitivity values a_i_ thereby determine its response to a given toxin dose. Fully sensitive cells (a_i_ = 1) are killed after only 1 hit (N_hits_= 1), but a_i_ < 1 (shields) may allow for N_hits_ > 1. Further details are provided in Fig. S1. **(C)** We simulate T6SS battles by randomly mixing cells on a 2-D surface. **(D)** Colormap charting positive selection (“Competitive advantage”; see colour bar) for rare mutants as a function of resistance to individual toxins (a_A_, a_B_), against attackers armed only with toxin “A” (red pac-men). **(E)** As (D) but against attackers armed with both “A” and “B” toxins (red and yellow pac-men). In D, E, colours show mean log-transformed Competitive advantage across N=20 10h replicate simulations, each initiated with random assortment of 500:90:10 (attacker:sensitive:resistant) cells. These data are replotted in Fig. S2 to show variances.

Figures 1B and S1 show an example of this model being used to represent two levels of resistance to an attacker bearing a single T6SS toxin “A” (red Pac-Men): a_A_ = 1 gives a maximally sensitive cell, requiring only a single T6SS hit to kill (N_hits_ = 1, Fig. 1B, left). If sensitivity a_A_ is reduced to 0.7, the target cell now requires 2 hits to kill it, and so has phenotypic resistance to that focal toxin (N_hits_ = 2, Fig. 1B, right). Our model allows us to consider a broad spectrum of possible sensitivities to T6SS toxins, ranging from a_i_ = 1 (one unit of toxin i is lethal) to a_i_ = 0 (total insensitivity to i). In this study, we focus primarily on comparisons between arsenals of 1 or 2 T6SS toxins (shown as red and yellow Pac-Men), but we also consider larger arsenals in the Supplementary Material (Fig. S4). We also assume throughout that toxin interactions are purely additive (toxin interaction coefficients B_ij_ = 0 for all toxins i,j), and that resistance mutations are free of fitness costs, but again test departures from these scenarios in the Supplement (Figs. S5 and S6 respectively).

### Simulated multi-toxin attackers constrain selection for resistance mutations

Against attackers armed with single or multiple T6SS toxins (“A” and “B”), when do we expect to see positive selection for resistant lineages? To address this question, we used our agent-based model (Fig. 1C) to simulate a simple selection scenario, asking whether a rare, cost-free mutant with reduced susceptibility to A and/or B would increase in frequency, in competition with populations of i) a given T6SS attacker and ii) fully susceptible cells. For a set of hypothetical resistance mutations spanning 0 ≤ a_A,_ a_B_ ≤ 1, we simulated tripartite competitions between populations of T6SS attackers (green), fully susceptible target cells (a_A,_ a_B_ = 1, magenta), and rare mutant target cells (a_A,_ a_B_ < 1, orange). We initiated simulations with a random arrangement of 500:90:10 attacker:susceptible:mutant cells, ran T6SS competitions for 10h (approximately 4 generations), and then calculated whether the focal mutant (a_A,_ a_B_) increased in frequency relative to its susceptible competitor, averaging over multiple simulation replicates (N=20 simulations per parameter combination) to account for model stochasticity.

Figs. 1D and S2A map selection for resistance mutants (measured using Competitive advantage, the final mutant:susceptible ratio divided by its starting value, see Methods), subjected to selection by attackers armed with toxin A only. These maps show that a wide range of possible resistance mutations are subject to positive selection (Competitive advantage > 1), including both small- and large-effect mutations, and mutations reducing sensitivity to single (a_A_ < 1, a_B_ = 1) or multiple toxins (a_A_, a_B_ < 1). As expected, the only mutations not subject to positive selection were those reducing sensitivity to the wrong toxin (a_B_ < 1, a_A_ = 1). In summary then, a wide range of possible rare resistance mutants are subject to positive selection. This suggests that, in the absence of resistance costs, resistance could readily evolve against a single-toxin attacker, assuming such sensitivity changes can arise via a single mutation.

In contrast, against a 2-toxin attacker, rare mutant selection was much more limited (Figs. 1E and S2B). Here, both small- and large-effect mutations, including those granting total immunity (a_i_ = 0) to one of the two toxins, were not subject to positive selection, even with no associated fitness cost. We intuit this result as follows: a mutant with protection against one toxin has no selective advantage over a fully susceptible competitor if simultaneously given a lethal dose of a second toxin: phenotypically, both succumb after the same number of T6SS hits. The only mutants subject to positive selection to 2-toxin attackers were those with substantially lower sensitivity to both toxins A and B. While such multi-resistance phenotypes can emerge from simple genetic changes (e.g. via alterations to exopolysaccharide synthesis regulation^27^), it seems unlikely that these adaptations are necessarily more likely than other small- and/or single-effect mutations. Subject to this assumption, this suggests that multi-toxin attackers will generally limit the evolution of T6SS resistance, since only mutations conferring strong multi-toxin resistance will be able to outcompete susceptible competitors.

### Simulations predict multi-toxin attackers limit stepwise evolution of T6SS resistance

Our model predicts that a major effect of T6SS toxin multiplicity is in constraining the set of non-costly resistance mutations subject to positive selection. So far, however, we have only considered resistance evolution as a single-step process; alternatively, resistance might evolve sequentially, via the accumulation of multiple small-effect mutations^40,46^. To investigate this possibility, we extended our model to consider multi-step evolutionary trajectories within the sensitivity parameter spaces shown in Figs. 1D and 1E. Based on our previous work in evolutionary game theory^43^, we devised a three-step loop to simulate evolutionary trajectories (Fig. 2A): beginning with a resident population of target bacteria with maximum sensitivity (a_A_, a_B_ = 1), we introduce a rare mutant with a random, cost-free perturbation to both its toxin sensitivities, (a_A_ + δa_A_, a_B_ + δa_B_). This perturbation represents a positive or negative change to each sensitivity parameter independently, arising from a single hypothetical mutation. If this mutant increases in frequency when subjected to T6SS selection pressure (simulated using our agent-based model), we assume that it will ultimately supplant and replace the resident – e. assuming i) no negative frequency-dependent selection^47^, and ii) a mutation rate low enough to avoid clonal interference^48^. If the mutant does not increase in frequency, then the resident is not replaced, and no evolutionary change occurs. A third possible outcome is that both the susceptible and the mutant population are driven extinct by attackers, in which case the evolutionary loop is terminated. In this way, we can predict the fates of sequences of small-effect mutations, and so assess potential for sequential resistance gain over evolutionary timescales.

**Fig. 2:**
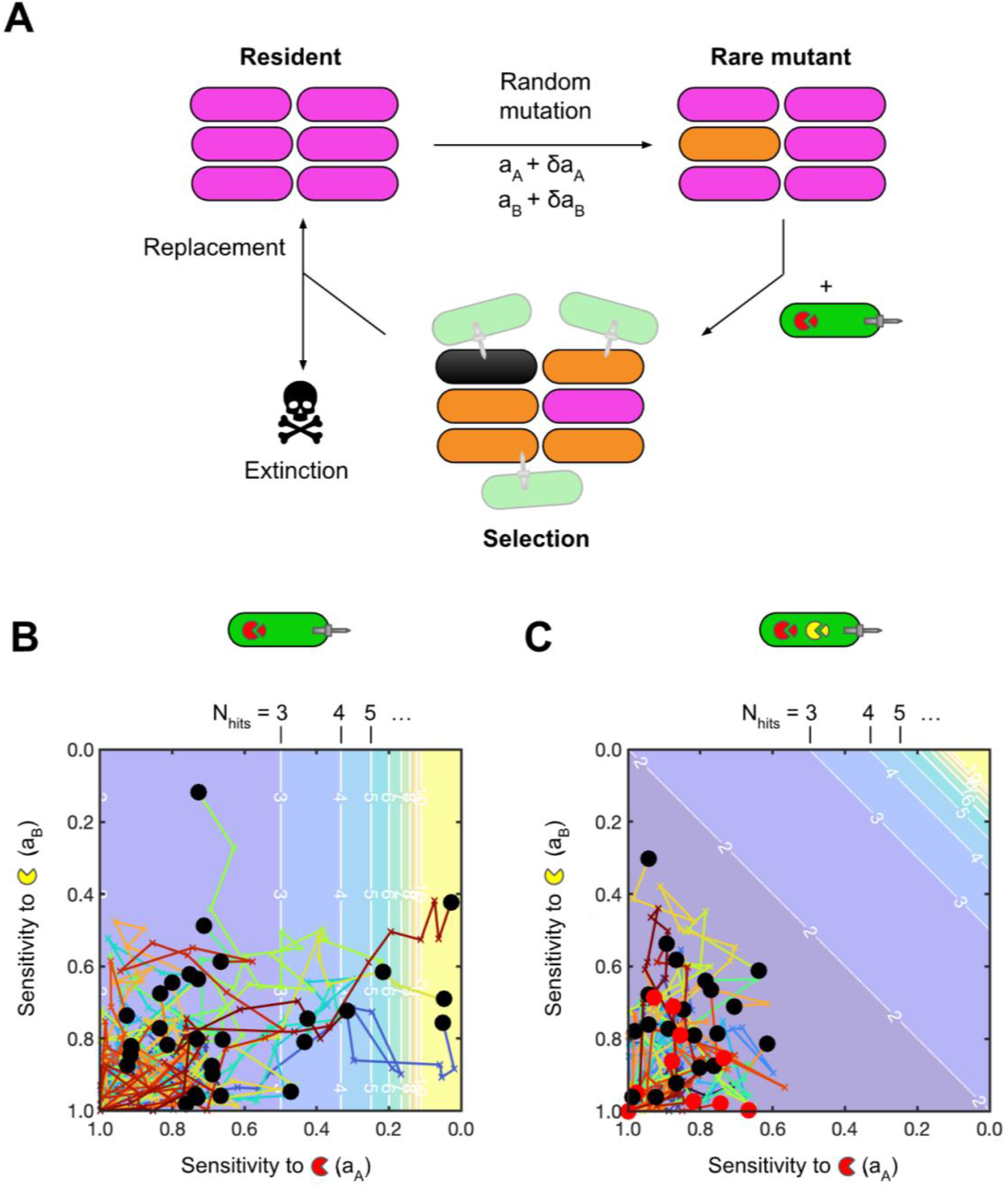
Multi-toxin attackers limit step-wise evolution of multi-toxin resistance. **(A)** Multi-step resistance evolution model, extending simulation results shown in Fig. 1. We use a 3-step loop to simulate evolutionary trajectories: i) a rare mutant is generated by randomly perturbing sensitivity levels to the two toxins A and B; ii) the resident and mutant populations are subjected to T6SS selection by a given attacker; iii) if the mutant increases in frequency relative to the resident, it replaces the resident. Total elimination of all target cells results in extinction and the loop is terminated. **(B)** Evolutionary trajectories against attackers armed only with toxin “A” (red pac-men). Each of N=30 trajectories (max. length 30 mutations) is shown as a single coloured line. Trajectories are plotted on a contour map showing the phenotypic resistance level (i.e. the number of T6SS intoxication events required to kill the focal cell, N_hits_). Black circles mark final sensitivity levels (a_A_, a_B_) final of resident population; red circles mark extinction events. **(C)** as with (B) but against attackers armed with both “A” and “B” toxins (red and yellow pac-men); N=30 trajectories. In (B,C), each simulation is initiated with a 500:90:10 mixture of attacker/resident/mutant cells, and lasts for 1.5h. All trajectories begin at (a_A_, a_B_) = (1,1). Mutations are drawn from normal distributions N(μ=0, σ=0.1); we consider other trajectory parameters in the Supplementary Material (Fig. S3).

In Fig. 2B and 2C, we plot evolutionary trajectories and their endpoints (black circles) generated with this three-step model, starting from a state of total susceptibility (a_A_ = a_B_ = 1), and again comparing attackers armed with a single toxin (Fig. 2B) vs. attackers with two toxins (Fig. 2C). Fig. 2B shows that, against an attacker secreting toxin A only, high levels of A-resistance frequently evolved: in N=30 trajectories spanning 30 consecutive mutations, all achieved at least double the starting resistance level (N_hits_ = 2; N_hits_ shown as a contour plot in the background of Fig. 2B), with 10% (3 / 30) exceeding N_hits_ = 20. Unexpectedly, we observed that trajectories often developed substantial resistance against toxin B as well, despite this being absent from the attacker’s T6SS arsenal. This unexpected gain of multi-resistance occurred through a combination of i) neutral drift (i.e. mutants with reduced a_B_ randomly increasing in frequency independent of selection), and ii) positive trait correlation (i.e. mutants with decreases in both a_A_, a_B_ increasing due to selection). In contrast: against a two-toxin attacker (Fig. 2C), we observed a different outcome: while all (N=30) trajectories displayed some movement away from the initial state of total susceptibility, no phenotypic resistance evolved (N_hits_ = 1 for all trajectory endpoints). Additionally, we observed that lineages would go extinct with greater frequency (10 / 30 trajectories, red circles in Fig. 2C). This suggested an additional barrier to resistance evolution: by being deadlier for a wider range of sensitivity values, multi-toxin attackers can drive lineages extinct before phenotypic resistance becomes fixed in a population. Overall, single-toxin attackers can generate surprisingly high levels of multi-toxin resistance via sequential resistance evolution, whereas multi-toxin attackers suppress resistance evolution entirely i) by limiting the pool of viable resistance mutations and by driving evolving lineages extinct.

To examine the robustness of our models’ predictions, we explored the effects of including other processes and factors in this evolutionary framework. First, we considered the effect of varying mutation effect size in a simplified version of the model: here, we replaced the (time-intensive) spatially explicit selection simulations with a simple analytical calculation of phenotypic resistance, assuming mutants become fixed if N_hits_(a_A_ + δa_A_, a_B_ + δa_B_)_mutant_ > N_hits_(a_A_, a_B_)_resident_ (see “Simplified Model”, Materials and Methods). Crucially, this simplified model recapitulated the findings of our agent-based simulations, but at a substantially reduced computational cost. Fig. S3 shows that there was a substantial window of mutation size σ in which multi-toxin attackers suppressed resistance evolution relative to single toxin attackers; the simplified model also recapitulated the previous prediction of single-toxin attackers driving multi-resistance evolution. Second, since T6SS-armed bacteria with >2 toxins appear to be common^49^, we repeated these analyses for attackers armed with 3 toxins, showing that larger toxin arsenals lead to enhanced resistance suppression (Fig. S4). Third, we studied the effect of introducing synergistic or antagonistic interactions between toxins^49^, which demonstrated that these interaction respectively further suppress or enhance resistance evolution against multi-toxin attackers (Fig. S5). Finally, we considered the inclusion of fitness penalties associated with T6SS resistance (Fig. S6), which have been demonstrated in previous work by MacGillivray *et al*^34^. Including a fitness reduction proportional to toxin insensitivity further suppressed resistance evolution against multi-toxin attackers, since multi-resistance then became inherently more costly than single-toxin resistance. While they differ in their construction and assumptions, these models agree that, qualitatively, multi-toxin attackers will suppress the evolution of resistance to the T6SS.

### Experimental evolution demonstrates increased extinction rate and resistance suppression by multi-toxin T6SS attackers

Our agent-based models yield a simple but powerful prediction: having a multi-toxin arsenal limits the evolution of bacterial resistance against the T6SS. Next, we sought to test this prediction empirically, using experimental evolution. Inspired by the recent work of MacGillivray and colleagues^34^, we conducted a serial transfer experiment tracking resistance evolution by *Escherichia coli* against *Acinetobacter baylyi* strains bearing a T6SS with either single or multiple toxins. Shown in Fig. 3A and Table S3, these strains comprise i) *A. baylyi* secreting a cell-wall-targeting amidase toxin (Tae1); ii) *A. baylyi* secreting a cell-membrane-targeting lipase toxin (Tle1); iii) *A. baylyi* secreting both Tae1 and Tle1 and iv) a T6SS-knockout *A. baylyi Δhcp* strain. Amidase and Lipase toxins are both highly abundant among T6SS-armed Proteobacteria (representing >80% of the 1,134 toxins reported by LaCourse *et al*.^49^) and so are a natural starting point for examining resistance evolution to individual toxins. Each *A. baylyi* strain has been characterised previously^12^; we additionally verified each strain’s genotype using whole-genome sequencing, and verified T6SS-dependent killing by performing a competition assay (Fig. 3B). In this assay, we co-cultured each attacker strain with the ancestral *E. coli* strain on agar for 3h (1:1 mixture of washed, density-normalised overnight cultures, see Materials and Methods) before recovering cells and using selective LB agar media to count surviving CFUs of each strain. This experiment confirmed that T6SS+ strains eliminate ∼99.9% of target *E. coli* under these conditions compared with our T6SS-negative *Δhcp* strain, producing a strong selective pressure for resistance. It also showed that the two-toxin attacker is more potent than either of the single-toxin attackers, showing ∼10x reduction in *E. coli* survival.

**Fig. 3:**
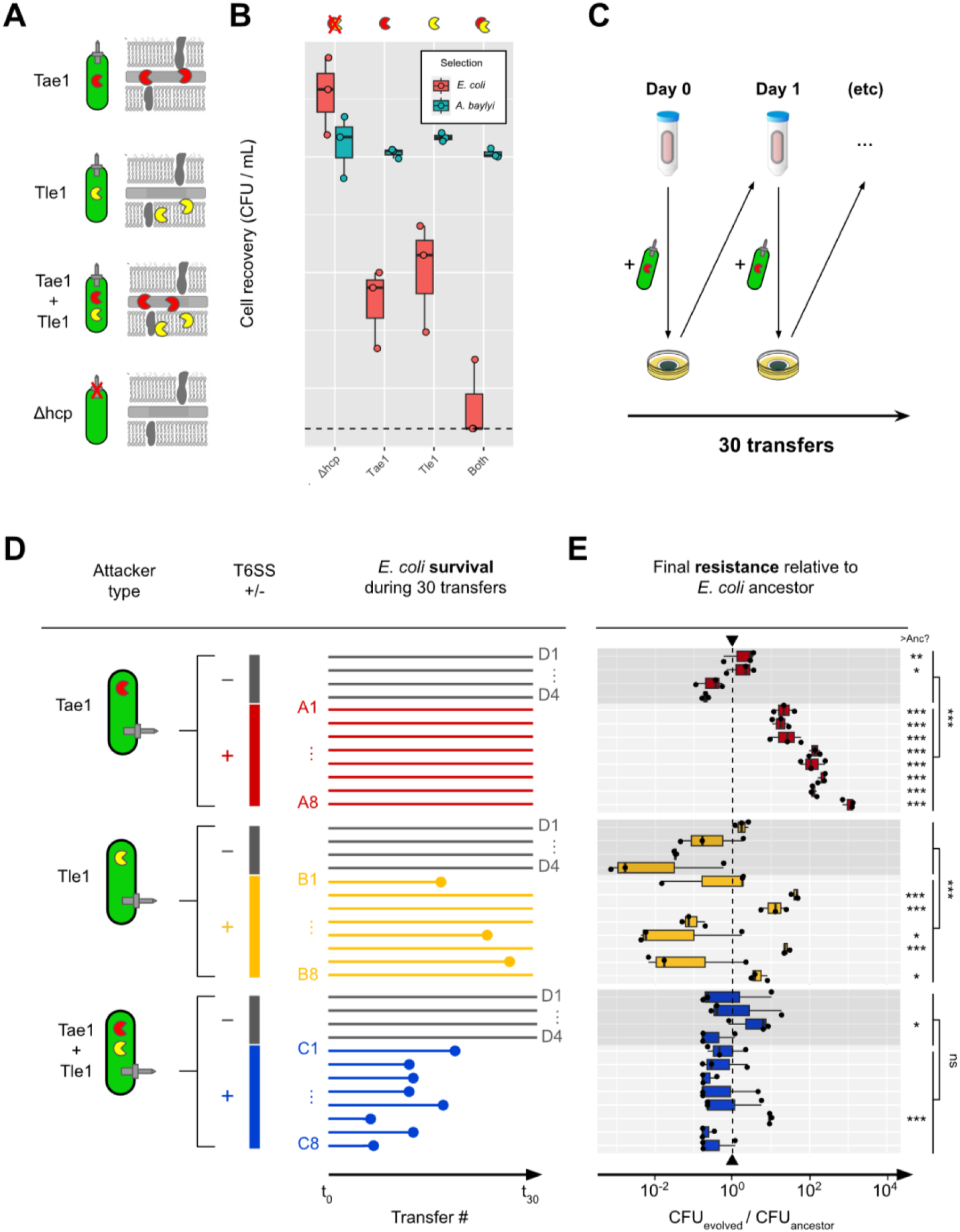
Experimental evolution demonstrates *E. coli* T6SS resistance suppression by multi-toxin *A. baylyi* attackers. **(A)** We evolved ancestral *E. coli* MG1655::*eGFP KanR* against attacker strains based on *A. baylyi* ADP1: these strains use their T6SSs to secrete amidase (Tae1), phospholipase (Tle1), or both toxins together (Tae1+Tle1). We also used T6SS- (*Δhcp*) ADP1 as a control for laboratory/coculture adaptation. **(B)** 1:1 competition assay comparing ancestral target (*E. coli*, pink) and attacker (*A. baylyi*, teal) cell recovery after coculture on nitrocellulose pads laid on agar (3h, 30°c), for each of the four attacker strains. **(C)** Experimental procedure for *E. coli* serial transfer (daily, in 3 blocks of 10 transfers). **(D)** Diagram comparing survival of each *E. coli* replicate line (coloured horizontal lines) between different T6SS attacker types. Extinction events are shown as filled circles. (**E)** End-point resistance of evolved lines against each focal treatment type, plotted relative to ancestral strain (dashed vertical line) and alongside *Δhcp*-evolved control lines (darker grey boxes); protocol as in (B). N=3 pseudobiological replicates (independent competitions blocked from same overnight cultures) shown in (B,E). Significance codes: ***, **, *, ns respectively denote increased survival relative to Anc, with p<0.001, p<0.01, p<0.05 and p≥0.05. Control- and treatment-evolved groups compared using ANOVA; individual isolates assessed for difference with ancestral strain using Dunnett’s test for multiple comparisons (applied to unnormalized survival data, Fig. S7).

Eight replicate lines of *E. coli* were subjected to 30 rounds of daily competition (∼500 generations) per T6SS+ attacker treatment, alongside 4 replicate control lines competed against the T6SS-negative *Δhcp* strain. Each serial transfer, we grew overnight cultures of each *E. coli* lineage in Kanamycin-supplemented Luria-Bertani (LB) broth, washing and mixing these with overnight LB cultures of the corresponding *A baylyi* attacker in a 1:5 ratio target:attacker ratio (again, ensuring a strong selective pressure for resistance). We then incubated these mixtures on pre-dried agar plates for 3h, before washing cells off each plate with a sterile buffer, and using the resulting cell suspension to inoculate the next day’s overnight culture. During the experiment, we monitored *E. coli* survival by taking daily optical density (OD_600nm_) readings of overnight cultures: lineages were deemed extinct if they showed no difference in optical density compared with media for 3 consecutive transfers. The three T6SS+ treatments showed marked differences in extinction rates (Fig 3D): Whereas all lineages survived until the end of the experiment in competition against the *Δhcp* control and Tae1 attacker, against the Tle1 attacker 3 / 8 lineages went extinct and against the double toxin attacker all lineages went extinct before the 20th transfer. This higher extinction rate is consistent both with the greater potency of the double toxin strain (Fig 3B) and with the predictions of our agent-based model (Fig. 2C): being more potent than single-toxin attackers, multi-toxin attackers were indeed more likely to drive susceptible populations extinct. Meanwhile, the intermediate extinction rate seen in Tle1-treated lines is consistent with Tle1’s higher potency under the conditions of the serial transfer experiment (Fig. S9,1:5 target:attacker ratio).

As well as driving extinction events, do multi-toxin attackers also suppress resistance evolution? To address this question, we isolated one endpoint clone from each of the 28 *E. coli* populations, at the latest sampling time point before extinction occurred (if it occurred). We then repeated our competition assays, subjecting washed overnight cultures of each endpoint clone to a 3h competition with its corresponding attacker strain, recovering cells, and measuring *A. bayly*i and *E. coli* survival. Fig. 3E plots the survival of each endpoint clone relative to the ancestral strain, highlighting the different levels of T6SS resistance that evolved in each treatment (raw data shown in Fig. S7). Against the Tae1 attacker strain, high levels of resistance evolved: in aggregate, these strains were ∼90x more resistant than the ancestor, and significantly more resistant than control lines D1-4 (ANOVA, p<2e-16). One isolate attained >3-log improvement in survival relative to the ancestral strain (A8, Dunnett’s test, p<0.001).

For lines evolved against the Tle1 attacker, outcomes were highly dependent on whether extinction had occurred. Isolates that avoided extinction (B2-4, B6 and B8) showed overall enhanced resistance (mean: 6x), with isolate B2 displaying a 43-fold increase in Tle1 resistance compared with the ancestral strain (Dunnett’s test, p<0.001). Of the isolates that went extinct, resistance levels were either unchanged (B1, B2, p>0.05) or significantly reduced (B5, p=0.0380) compared with the ancestral strain, suggesting that only surviving lines were subject to evolutionary rescue^50^. Finally, lineages evolved against double toxin attackers showed no significant resistance relative to the ancestor (mean: 0.6x) and control lines (ANOVA p=0.109), with the exception of isolate C6 (∼10x higher relative survival). Finally, for each T6SS attacker, *Δhcp* control isolates (D1-4) generally did not show significant increases in resistance (exceptions: D1 and D2 vs. Tae1 and D3 against Tae1+Tle1), suggesting that improved *E. coli* recovery in evolved isolates can be attributed to T6SS resistance proper, rather than generic adaptation to our lab conditions.

### Single-toxin T6SS attacks can select for cross-resistance to other toxins, driving multi-resistance evolution

Together these experimental data confirm two key predictions of our agent-based and mathematical models, namely that multi-toxin attackers would i) result in more frequent extinction events, and ii) suppress the evolution of T6SS resistance. Another prediction of our model was that single-toxin attackers could, counter-intuitively, result in the evolution of multi-toxin resistance. To test this prediction, we tested survival of each resistant endpoint clone against each of the four *A. baylyi* attacker strains (Figs. 4A, 4B and S8). We observed asymmetric cross-resistance relationships between Tae1-evolved and Tle1-evolved isolates (Fig. 4A). While Tle1-evolved isolates tended to show unaltered (B3, p>0.05) or increased (B4, B6 and B8, p<0.01) sensitivity to the alternate toxin Tae1 (overall: 1.76x reduction in resistance relative to ancestor), Tae1-evolved isolates tended to be cross-resistant to the alternate toxin Tle1 (A1, A3, A5, A7 and A8; p<0.05; overall 12.5x increase in resistance). In contrast, we found that isolate C6, having previously showed only marginal resistance to the two-toxin attacker, also showed no significant resistance to either Tae1 or Tle1 alone (p>0.05).

**Fig. 4:**
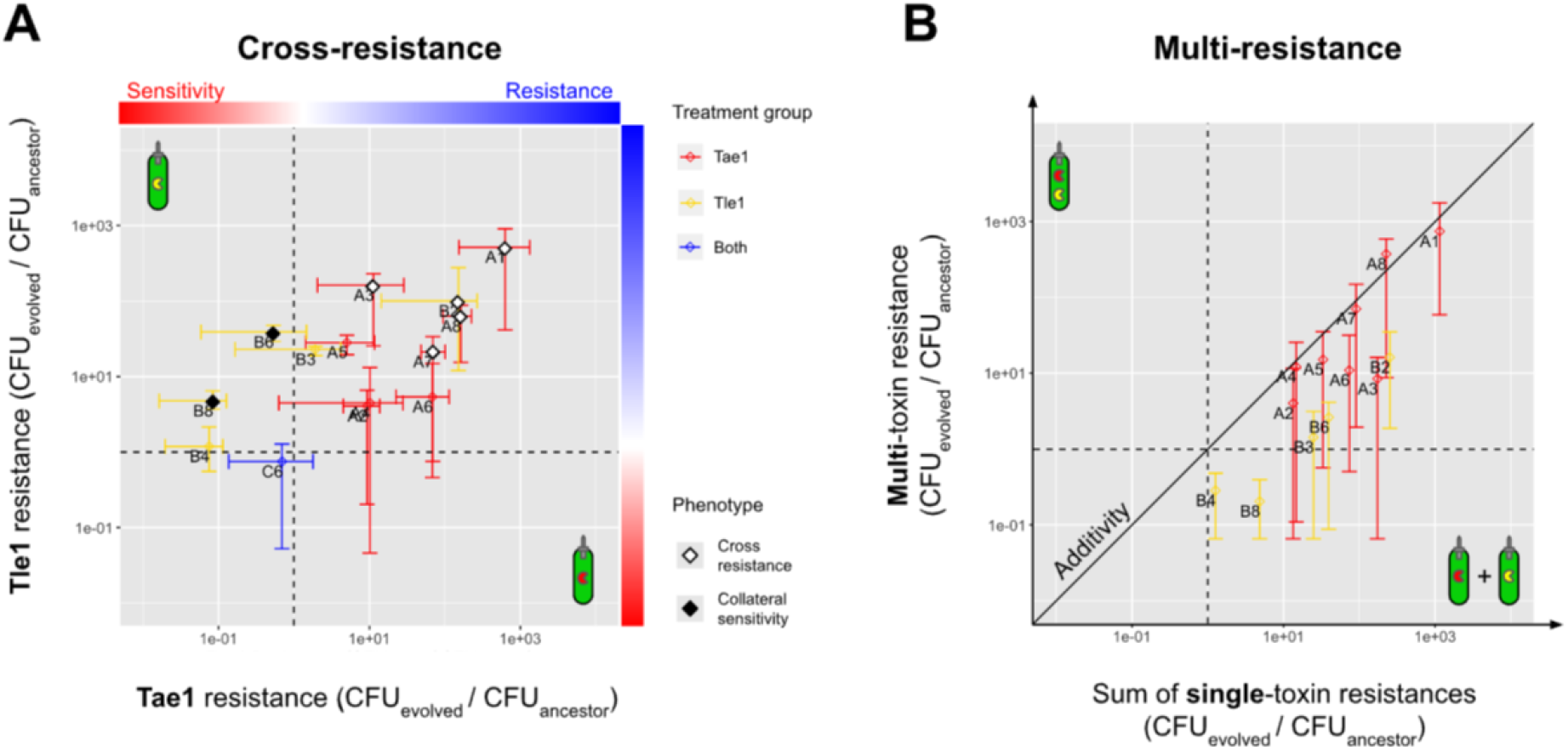
*E. coli* evolved against single-toxin attackers show cross- and multi-resistance, depending on treatment. **(A)** Factorial resistance assay showing relationships between Tae1 and Tle1 resistance (relative to ancestral strain) for a subset of evolved isolates, highlighting examples of cross-resistance and cross-sensitivity (white and black diamonds, respectively indicating i) significant increases to both focal *and* non-focal toxin; significant increase in focal toxin resistance *but* significant reduction in non-focal toxin resistance). Points and crosshairs indicate data means and ranges, coloured according to the toxin treatment against which that isolate evolved. **(B)** Relationships between mean resistance to individual toxin treatments (Tae1 resistance + Tle1 resistance, horizontal axis) and resistance to multi-toxin treatment; points coloured as in (A) Points and bars denote survival mean and range vs. the two-toxin attacker. Isolates with significant cross-resistance (gain of resistance to both focal and non-focal toxin) and cross-sensitivity (gain of resistance to focal toxin but loss of resistance to non-focal toxin) are respectively marked with white and black diamonds. N=3 pseudobiological replicates (independent competitions blocked from same overnight cultures) shown in (A, B). Individual isolates assessed for difference with ancestral strain using linear modelling (Dunnett’s test for multiple comparisons); unnormalized survival data and statistical tests shown in Fig. S8.

Moreover, the cross-resistance phenotype seen in A lines was also predictive of multi-resistance (Fig. 4B): resistance against the double-toxin attacker positively correlated with the sum of resistances against each single-toxin attacker alone. Isolates A1, A7 and A8 all displayed significant resistance to the two-toxin attacker *Ab*Tae1Tle1 (Fig. S8; p<0.05), consistent with their strong resistance to each of Tae1 and Tle1 individually. Finally, to determine whether these phenotypes were the result of T6SS resistance *per se* (and not, for instance, evolved increases in *E. coli* growth rate or similar), we also enumerated strains’ recovery against the T6SS-negative control *AbΔhcp* (Fig. S8B). This experimental control demonstrated that, while some evolved isolates (A1-3, A5, A7-A8 and B2) demonstrated ∼10x greater recovery in the absence of T6SS killing, these effects alone are insufficient to explain the resistance strengths observed in our resistance assays. We therefore conclude that the enhanced *E. coli* recovery seen in Figs. 3, 4, S7 and S8 are the result, at least in part, of T6SS resistance proper.

Overall, these data support our models’ two main predictions: i) that multi-toxin attackers can suppress resistance evolution, and ii) that single-toxin treatments can, counterintuitively, promote the evolution of multi-toxin T6SS resistance.

## Discussion

When it comes to interbacterial warfare, are two toxins better than one? The ability to inject multiple toxins at once is a rare feature among bacterial weapons^18^. In contrast, T6SS-armed bacteria often carry multiple toxins: of 474 Proteobacterial species surveyed by LaCourse *et al*, approximately 40% carried >1 functionally-distinct antibacterial effector^49^. The potential advantages of having a broader toxin arsenal are several: compared with single toxin analogues, multi-toxin attackers can target a broader set of competitor strains and species, over a wide range of environmental conditions. Aside from having increased overall potency, due to the additive toxicities of their toxins, multi-toxin attackers may also benefit from toxin synergies, although toxins can also display anti-synergistic interactions^49^.

Our work reveals an additional benefit to having a functionally diverse T6SS toxin arsenal: resistance evolution suppression. Using agent-based models, we predicted that bacteria using multi-toxin T6SS attacks could suppress resistance evolution against this widespread and ecologically significant antibacterial weapon. This suppression occurred via two distinct mechanisms: multi-toxin attackers i) constrained the set of mutations subject to positive selection, and ii) drove target populations extinct more frequently, precluding further adaptation. Our evolutionary experiments confirmed these predictions, demonstrating multi-toxin attackers’ greater potency and commensurate ability to drive target populations extinct, and so limit resistance evolution. Of course, this resistance suppression may simply be an incidental by-product of having multiple toxins, rather than an evolved function *per se*. However, the ability to maintain lethal anti-competitor weaponry is clearly important for the success of many bacteria^25,51^, and so conceivably resistance suppression could partly explain the abundance of multi-toxin arsenals across T6SS-weiling bacteria^49^.

We also found that some single-toxin T6SS attackers were, surprisingly, more likely to generate multi-toxin resistance than multi-toxin attacks, mirroring similar results on resistance to antimicrobial combinations by Gifford *et al*.^46^ Our modelling suggests that this process could occur via several mechanisms: i) through positive genetic correlation, where non-focal resistance is selected for because it also grants resistance to the focal toxin; or ii) through drift, where non-focal resistance becomes fixed at random, especially where associated fitness costs are low. In this framework, however, it remains unclear why our single-toxin Tle1 treatments selected largely for collateral sensitivity instead of cross-resistance. Our Tae1-adapted populations show strong resistance to Tle1, so why do these same adaptations not appear in our Tle1-adapted populations? In any case, further work is needed to characterise the genetic and molecular bases of our resistant *E. coli* isolates, and the evolutionary dynamics through which they emerge and establish^52^.

Our computational and experimental approaches both have several limitations. First: while biophysically detailed, our models still present a highly simplified view of T6SS-mediated competition, ignoring various environmental and ecological details (e.g. solute gradients, within-population heterogeneity, macroscale spatial structure) that may be present within specific natural communities. Second, our stepwise evolution model assumes that resistance mutations are rare, such that they are subjected to selection in isolation. This neglects the potential role of clonal interference, which might present an additional barrier to resistant mutant fixation. Third, our models focus on individual level traits, neglecting the potential for collective resistance mechanisms^27,28^. While our experimental parameters are designed to select for resistance at the single cell level (by using high cell density and high attacker:target cell ratio, so that a maximum number of target cells are individually subject to T6SS attack^34^, we cannot say which of our evolved isolates have a collective component to their T6SS resistance. Clearly both individual- and collective level defences exist against the T6SS^22^, and an interesting avenue of future research would be to compare the circumstances that lead to one form of resistance or another.

The ability of multi-toxin attackers to limit resistance evolution has an obvious technological attraction. This property could recommend the multi-toxin T6SS weaponry as a basis for future biocontrol agents^53^, or as live antimicrobial biotherapeutics^54,55^. Naturally, claims of resistance evolution suppression should be treated with caution^56^, particularly when discussing potential new treatment strategies. This is underscored by the fact that multiple mechanisms of T6SS multi-resistance are already known to exist^20,22^: these include biofilms and capsule formation^27,31^, which appear to physically block T6SS injection and thereby prevent any toxin translocation^30^. Multi-resistance against T6SS-armed *V. cholerae* bacteria has also been shown to evolve spontaneously via several independent mechanisms^34^. Just as existing clinical forms of combination therapy are imperfect in suppressing resistance^38^, we should not expect T6SS-based antimicrobials to be “resistance-proof”. Nevertheless, even imperfect resistance suppression is a desirable property for an antimicrobial, and this may make T6SS-based biocontrol and biotherapeutic strategies a more attractive future prospect.

The T6SS is a widespread and ecologically significant antimicrobial weapon. Simultaneous multi-toxin delivery—rare among other antimicrobial weapons—is an important but poorly-understood feature of the T6SS, stemming naturally from its contractile mode of action. Our work shows that, analogous to clinical combination therapies, multi-toxin arsenals limit resistance evolution to bacterial T6SSs, shedding light on the selective forces shaping microbial attack and defence evolution, and helping to explain why many T6SS-armed bacteria encode multiple toxins. The inherent ability to limit resistance evolution is a highly desirable feature in an antimicrobial; our findings may recommend the T6SS for future applications as an effective biotherapeutic or biocontrol agent.

## Materials and Methods

### Agent-based modelling

#### Software code

We used the open-source modelling software *CellModeller*^41^ to simulate the evolution of T6SS resistance, in combination with custom Python modules published previously^27^. These modules are available for download at https://github.com/WilliamPJSmith/CellModeller). Below, we provide a detailed description of our model and the alterations made to it in this study. Model variables and parameters are summarised in Tables S1 and S2 respectively.

#### Cell growth and division

We model bacterial cells as elongating spherocylindrical rods, having fixed radius R = 0.5μm and variable segment length L. Each cell’s volume V increases exponentially with specific growth rate k_grow_ as dV / dt = k_grow_ V until it has approximately doubled its birth volume V_0_; upon reaching volume V = 2 V_0_ + η (with η representing uniform random noise in division volume), the cell divides lengthwise into two identical daughter cells. Repulsive elastic forces between cells are simulated by completing an energy minimisation algorithm at each simulation timestep, as described previously^57^. This process computes minimal cell translations and rotations to iteratively remove cell-cell overlap caused by growth.

#### T6SS firing and toxin doses

Every simulation timestep, attacker-type cells fire N_firing_ times, with N_firing_ drawn for each cell individually from a Poisson distribution with mean k_fire_ Δt, with k_fire_ the population firing rate (units: firings cell^-1^ h^-1^) and Δt the simulation timestep (h). Attacker-type cells fire T6SS needles from randomly chosen points on their cell membranes; each needle has a length of one cell radius, R=0.5μm. After firing, all needles and cells are sorted according to their location in the x-y plane using an infinite square grid of element size 10μm; a line segment hit detection algorithm is applied to compute which needles strike which cells^42^. Successful hits translocate specific doses d_i_ of individual toxins to the first cell struck by the focal needle, depending on the attackers’ cell type: in this study, single-toxin attackers translocate unit doses d_i_ of toxin “A” only (d_A_ = 1, d_B_ = 0); two-toxin attackers translocate one of each toxin (d_A_ = 1, d_B_ = 1).

#### T6SS toxin responses and resistance

In contrast to our previous models^27,42,43^, we allow cells to respond to toxin translocation in a step-wise, dose-dependent manner, as follows. Each cell has a scaled integrity variable I set to I=1 at birth. Individual toxins damage cells, depleting their integrity in proportion to their cumulative intracellular concentration x_i_ (units: toxins cell^-1^), weighted by a sensitivity parameter a_i_; the overall damage (i.e. loss of integrity) following a T6SS translocation event is assumed to be the sum of damage terms over all N_toxins_ toxins. To account for possible toxin synergy, antagonism or non-linear responses^49^, we also include a second term representing pairwise interactions between toxins, which can enhance or mitigate their toxicity according to the sign and magnitude of toxin interaction parameters B_i,j_. The overall integrity equation is then

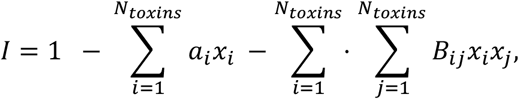

where a_i_, B_i,j_, and x_i_ are respectively toxin sensitivity parameters, toxin interaction parameters, and cumulative intracellular concentrations for toxins i and j. Cells are killed (setting k_grow_ = 0) once their integrity falls below zero, and subsequently lyse (are physically removed from the simulation) after a set time delay, 1 / k_lysis_. Toxin activity is assumed to be constant over the course of a simulation, such that it is necessary only to know a cell’s accumulated dose of each toxin type {x_i_} to determine whether it has been killed. In our main text, we set B_i,j_ = 0 (purely additive interactions), but we consider scenarios in which |B_i,j_| > 0 (non-additive interactions) in the supplement (Fig. S5). With B_i,j_ = 0, a cell’s set of sensitivity parameters, {a_i_}, controls its sensitivity to each individual toxin, and determines how many translocation events it can sustain from a particular attacker (N_hits_), according to the equation

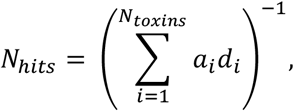

where d_i_ is the dose of toxin i translocated per successful T6SS hit (dependent on attacker type), and a_i_ is the sensitivity to toxin i (dependent on target cell type). Attackers are assumed to be perfectly immune to their own complement of T6SS toxins (a_i_ = 0 for each toxin i carried), and so have N_hits_ = ∞ against their kin.

While simplistic, this model has several useful features. First, it allows us to independently vary sensitivity to an arbitrary number of individual T6SS toxins, and so compute how many T6SS translocation events a cell can withstand (phenotypic resistance, N_hits_) as a function of the toxin(s) being secreted by attacker cells, and the target cell’s sensitivity to each. Second, it also can represent cross-resistance (and cross-sensitivity) phenotypes^58^, i.e. where changes in one sensitivity parameter a_i_ are positively (negatively) correlated with changes in sensitivities to other toxins. Third, it enables us to parameterize positive and negative interactions between toxins (via interaction terms B_ij_), capturing potential non-additive interactions^49^ between toxins, and non-linear dose responses (by setting |B_ii_| > 0).

#### Simulation parameters

In our initial resistance selection simulations (Figs. 1, S1 and S2), we initialised simulations with a mixture of attacker (a_A_ = a_B_ = 0), susceptible (a_A_ = a_B_ = 1) and mutant (a_A_,a_B_ variable) cells in the ratio 500:90:10, such that i) susceptible cells would face strong selective pressure from attackers, and ii) the mutant began at low frequency relative to the susceptible strain. Cells were initially placed and orientated randomly on a flat surface, within a circular area of radius 50μm. We then allowed cells to grow, divide and compete for a period of 10h (approximately 4 doubling events, chosen to balance selection effect size against computational speed), whereafter simulations were terminated and the initial and final frequencies of each cell type compared. We used these to compute the Competitive advantage of mutant strains relative to the more common susceptible strain, using the equation

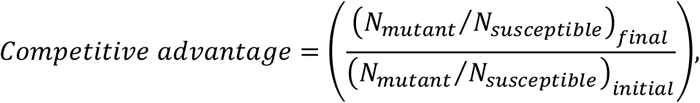

where N represents the population size of the mutant or susceptible strain at the indicated (initial or final) timepoint.

#### Evolutionary trajectory simulations

To model step-wise resistance evolution (Figs. 2, S3-S6), we ran chains of simulations sequentially, simulating the stepwise evolution of resistance via the accumulation of multiple small-effect mutations. We began with a 500:90:10 mixture of attackers, “resident” sensitive cells (a_A_ = a_B_ = 1 initially) and “mutant” sensitive cells with some perturbation δa_i_ to the resident strain’s sensitivity, (a_A_ + δa_A_, a_B_ + δa_B_). Each mutation δa_i_ was drawn independently from a normal distribution with mean 0.0 and variance 0.01 (i.e., allowing mutations that independently alter sensitivity to multiple toxins); mutations resulting in sensitivities outside of the range 0 ≤ a_i_ ≤ 1 were automatically rejected and re-drawn. Simulations were initialised as above and terminated after 1.5h. If focal mutant strain increased in frequency relative to the resident in one simulation, then it replaced the resident strain in the next simulation, which was then initialised and run as before. We repeated this process to generate trajectories of N=30 simulations (Fig. 2), as well as longer trajectories for our simplified fitness model (described below).

#### Computation

Agent-based simulations were run using i) a 2014 iMac desktop computer (3.5 Ghz Quad-core Intel i5 core) and ii) using the University of Manchester’s Computational Shared Facility (CSF3, MERMan-dedicated compute nodes), using pairs of *NVIDIA*^*TM*^ v100 Volta GPUs.

### Simplified fitness model

While incorporating many biological details of T6SS battles, our agent-based model is nonetheless computationally demanding, requiring multiple days of computation time to simulate evolutionary trajectories. To permit a broader range of robustness tests, we created a simplified version of our agent-based evolution model (Fig. S3A), altering the trajectory loop structure so that the test for resident replacement is replaced with a simple comparison of phenotypic resistance,

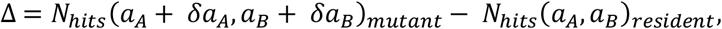

where, as above,

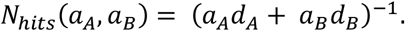

If the mutant strain’s T6SS resistance exceeds that of the resident (Δ>0), it replaces it as in the original trajectory model. This simplification greatly accelerates trajectory simulation, at the cost of removing key ecological processes featured in the agent-based model (particularly: community spatial structure, frequency-dependent fitness, stochasticity and extinction events). Two key consequences of these omissions are i) lineage extinction is no longer possible, but ii) the barrier to mutant fixation is greater in the simplified model, since neutral or near-neutral mutations can no longer become fixed through drift. In practice however, the agent-based and simplified trajectory models agree qualitatively that single-toxin attackers facilitate more single- and multi-toxin resistance evolution than do multi-toxin attackers (Figs. 2B and S3B).

#### Computation

We simulated evolutionary trajectories (Figs. S3-S6) with the simplified fitness model using *Matlab* version 2023b, running on a 2023 Apple MacBook Pro (16”, M2 Chipset) laptop computer.

### Experimental evolution

#### Bacterial strains

Table S3 summarises bacterial strains used in this study. We used whole-genome short-read sequencing (*MicrobesNG*^*TM*^ standard Illumina sequencing service, 250 bp paired-end reads at 30x coverage) to confirm the identity of each strain.

#### Establishing E. coli populations

8 + 8 + 8 + 4 = 28 independent populations of *E. coli* AncG were established by streaking 25% glycerol cryostocks onto 1.5% w:v LB agar plates, incubating overnight (∼18h) at 30°C until individual colonies became visible. Individual colonies were then selected at random and used to inoculate 28 identical cultures (polypropylene tubes each containing 5mL LB + 25μg/mL Kanamycin sulfate) using sterile plastic loops. An identical tube was also sham-inoculated (loop but no colony); this “sentinel” tube acted as an internal contamination control for the duration of the experiment. All cultures were then incubated overnight (Eppendorf New Brunswick™ Innova 44 shaking incubator; 180RPM orbital shaking at 37°C) with lids taped on loose (¼ turn) to ensure aeration.

#### Attacker cultures

Fresh ADP1 cell cultures were grown daily by inoculating cryostocks directly into 5mL LB as above, incubating overnight (180RPM orbital shaking at 30°C).

#### Daily Passaging

As in previous work^34^, *E. coil* cultures were subjected to 30 daily rounds of competition, in this case with one of four *A. baylyi* attacker types: Tae1 (8 replicates), AbTle1 (8 replicates) or AbTae1Tle1 (8 replicates). Control *E. coli* cultures were passaged against T6SS-negative Ab*Δhcp* (4 replicates), to examine adaptation to *A. baylyi* in the absence of T6SS activity, and to lab culture conditions. In each round of competition, we washed 100μL of each EC overnight culture once with an equal volume of fresh LB to remove antibiotics (4 mins at 20,000g, ThermoFisher Scientific Heraeus Pico 17 benchtop microcentrifuge). Similarly, 5mL of each ADP1 overnight culture was washed once with 5mL fresh LB, using gentle centrifugation (3,500RPM, 10 minutes, Sigma 2-6E benchtop centrifuge). Washed ADP1 and EC cultures were then pipette-mixed at a 5:1 attacker:defender ratio, and 50μL of each mixture was spotted onto the centre of a pre-dried, pre-warmed 60mm LB agar plate, dried for 20 minutes, and then incubated at 30°C for 3 hours. Following incubation, cells were recovered and resuspended by repeatedly jetting spots with 1mL sterile dPBS; 500μL of this recovery mixture was then transferred into 5mL fresh LB supplemented with 25μg/mL Kanamycin sulfate, inoculate the next round of overnight cultures. To monitor overnight growth during serial passaging, the optical density (OD at 600nm, OD_600_) of 10x diluted *E. coli* and *A. baylyi* overnight cultures was measured using a Clariostar Plus (BMG LabTech) or SPARK multimode (Tecan) plate reader. *E. coli* cultures that failed to become turbid overnight for three consecutive nights were designated extinct and withdrawn from the passaging regime.

#### Batches and sampling

Every 5 transfers, 500μL samples of each overnight population were mixed with 500mL 50% v:v sterile glycerol and frozen at -80°C. We repeated the transfer procedure daily for 10 rounds, whereafter each population was frozen as above to ‘pause’ the experiment. Whole-population samples were then used to restart the next 10 transfers, thawing cryotubes for 45 mins at room temperature before transferring 500mL thawed stock directly into Kanamycin-supplemented LB as before, to inoculate the next round of overnight cultures. We performed 3 blocks of 10 transfers in this way, for a total of 30 daily transfers. To ensure that each replicate received the same treatment on average, the order in which treatments and replicates were processed was reversed every 5 transfers. Following the completion of the 30 transfers, we isolated clones from the last surviving sample taken from each cell line, streaking populations on 1.5% w/v LB agar, and picking colonies as before. These end-point clones were then grown overnight in 5mL LB, frozen as 25% v:v cryostocks, and later subjected to further phenotypic testing, as detailed below.

### Initial resistance assay

#### Competition setup

Based on previous work^12^, we conducted a competition assay to measure each evolved isolate’s resistance to its corresponding treatment type (Figs. 3E and S7), relative to the (anc)estral *E. coli* strain (Fig. 3B). As above, we grew 5mL LB overnight cultures of each end-point *E. coli* clone (28 isolates) and each *A. baylyi* attacker (4 strains), washing cultures once in an equivalent volume of fresh LB (500μL for *E. coli*; 5000μL for *A. baylyi*), and then normalising densities to OD_600_ = 1.9 for *E. coli*, or 0.9 for *A. baylyi*. These densities reflect the average densities of overnight *E. coli* and *A. baylyi* cultures, and so allow us to replicate the physiological conditions of our evolutionary transfer experiment in a fair and reproducible manner. We then combined the washed and normalised *E. coli* and *A. baylyi* cultures in a 1:1 ratio (90μL + 90μL), using pipette-mixing to homogenise mixtures, as follows:

1. *E. coli* isolates Anc, A1-8 and D1-4 with *Ab*Tae1;
2. *E. coli* isolates Anc, B1-8 and D1-4 with *Ab*Tle1;
3. *E. coli* isolates Anc C1-8 and D1-4 with *Ab*Tae1Tle1;
4. *E. coli* isolate Anc with *AbΔhcp*.

We plated 50μL of each co-culture mixture in triplicate on sterile 13mm Mixed Cellulose Ester (MCE) filter papers (pore size 0.22μm, Merck Life Science UK Ltd.), placed flat on pre-dried, pre-warmed agar petri dishes, each filled with 5mL of 60mm 1.5% w/v LB agar. Each co-culture droplet was allowed to dry completely in a laminar flow cabinet, before plates were lidded, inverted and transferred to a static incubator (3h, 30°c). To investigate the potency of each T6SS treatment vs. the Ancestral *E. coli* strain, we also repeated this experiment using the 1:5 target:attacker ratio used during our serial transfer experiment (Fig. S9).

#### Cell recovery and Colony-Forming Unit (CFU) counting

After incubation, we recovered cells by transferring each MCE filter into a 15mL falcon tube containing 1mL of sterile dPBS, and then vortexing the tube for 10s to recover surviving cells. We used this cell suspension to make a dilution series (7x 10-fold dilution in dPBS), plating 5μL of each dilution in triplicate on pre-dried, pre-warmed 120mm square vented petri dishes filled with 40mL 1.5% w/v LB agar plates, supplemented either with 25μm / mL Kanamycin sulfate (selecting for *E. coli*) or 100μm / mL Streptomycin sulfate (selecting for *A. baylyi*)^12^. These plates were then incubated at either 37°c (*E. coli*) or 30°c (*A. baylyi*) overnight (∼16h) until individual colonies became visible. To enumerate each strain’s survival, we counted the number of CFUs at the highest dilution at which at least 3 CFUs were visible, and then multiplied this number by the dilution factor. To account for plate-to-plate variation in CFU count from a given recovered cell suspension, we took the mean count over 3 replicate platings for each competition replicate.

### Combinatorial cross-resistance assay

To compare evolved isolates’ resistance to different T6SS toxins, we repeated our initial resistance assay with a subset of evolved isolates (Figs. 4 and S8): specifically, those that survived the entire 30-transfer experiment (A1-8, B2-4, B6, B8; see Fig. 3D). We also included isolate C6, which, despite surviving only 19 transfers, appeared to have developed weak resistance to *Ab*Tae1Tle1 in our initial screen. As a test of resistances’ dependence on growth phase, we began this experiment with exponential-phase cultures of each culture, as follows: 5mL LB shaking cultures of the above *E. coli* isolates, ancestral *E. coli* strain, and *A. baylyi Ab*Tae1, *Ab*Tle1, *Ab*Tae1Tle1 and *AbΔhcp* strains, were grown overnight as above. Respectively, 100μL (1:50 dilution) of *E. coli* overnight cultures, and 1000μL (1:5 dilution) of *A. baylyi* overnight cultures, were transferred into 5mL fresh LB and incubated in a shaking incubator (as before: 180RPM orbital shaking at 37°C for *E. coli*, 180RPM orbital shaking at 30°C for *A. baylyi*) for 2.5h, until cultures reached an OD_600_ of ∼1.0. We then washed cultures once in equivalent volumes of fresh LB as above, normalised optical densities to OD_600_ = 0.1, and pipette-mixed cultures in a 1:1 ratio (100μL + 100μL) as follows:

1. *E. coli* isolates Anc, A1-8, B2-4, B6, B8, C1 with *Ab*Tae1;
2. *E. coli* isolates Anc, A1-8, B2-4, B6, B8, C1 with *Ab*Tle1;
3. *E. coli* isolates Anc, A1-8, B2-4, B6, B8, C1 with *Ab*Tae1Tle1;
4. *E. coli* isolates Anc, A1-8, B2-4, B6, B8, C1 with *AbΔhcp*.

These mixtures were then plated onto MCE filters on LB agar, dried, inverted and incubated for 3h. We then used the same protocol as above to recover and enumerate *E. coli* survival.

### Data processing and statistics

Agent-based model simulations were visualised using *Paraview* software version 5.4.0 and *Matlab* version 2023b. Data were analysed and visualised using *RStudio* version 2023.12.1+402, using the packages *readxl, ggplot2, cowplot, dplyr, tidyr* and *multcomp*, and using the *Matlab redblue* package (© 2009, Adam Auton). To compare levels of resistance between control (D1-4) and treatment (A-, B- or C1-8) isolates, we log10-transformed survival data (CFU / mL) to homogenize variances, and then used linear modelling with ANOVA (via R’s inbuilt *lm* method) to compare control vs. treatment, both normalised by ancestral survival. To compare individual isolates with the ancestral strain, we used Dunnett’s test for multiple comparisons (*multcomp* methods: *glht, mcp*), applied to unnormalized data (Figs. S7, S8).

## Contributions

WPJS, MB and MAB designed the project and secured funding. WPJS designed and implemented the model with EAB, and analysed the data with EAB and KZC. WPJS performed the experiments and analysed the data with CGK, MAB and MB. WPJS and MAB wrote the manuscript with feedback from EAB, KZC, CGK and MAB.

## Acknowledgements

We would like to thank Rosanna Wright, Anastasia Kottara, Matt Shepherd, Anna Richmond and other members of the Manchester Microbial Eco-Evolutionary Research (MERMan) group for vital equipment lends, technical support and wet lab training. We are indebted to Elisa Granato, Lin Lin, Alejandro Tejada Arranz and Mike Bottery for scientific consultation throughout the project, and to Claudia Igler for her comments on the manuscript. WPJS is funded by a Sir Henry Wellcome Postdoctoral fellowship award, 222795/Z/21/Z. KZC is supported by a Wellcome Trust Career Development Award, 226047/Z/22/Z.

