## Supplementary Information for "Multiplicity of Type 6 Secretion System toxins limits the evolution of resistance"

**Supplementary Tables**

**Table S1: Agent-based model variables**

| **Variable for cell *i*:** | **Symbol** | **Units** |
|---|---|---|
| Position vector | $\boldsymbol{p}_i = (p_x, p_y, p_z)_i$ | µm |
| Orientation unit vector | $\boldsymbol{v}_i = (v_x, v_y, v_z)_i$ | - |
| Segment length | $L_i$ | µm |
| Volume | $V_i = 4\pi R^3/3 + \pi L_i R^2$ | $\mu m^3$ |
| Intracellular toxin concentrations | $\boldsymbol{x} = (x_A, x_B)$ | $cell^{-1}$ |
| Cell integrity | $I = 1 - a_A x_A - a_B x_B - B_{AA} x_A^2 - B_{BB} x_B^2 - B_{AB} x_A x_B$ | - |

**Table S2: Agent-based model parameters**

| Type | Parameter | Symbol | Value(s) | Units | Source |
|---|---|---|---|---|---|
| **Cells** | Cell radius | $R$ | 0.5 | μm | Rudge2011 |
| | Cell volume at birth | $V_0$ | 0.54 | $\mu m^3$ | Smith2017 |
| | Max cell growth rate | $k_{max}$ | 1.0 | $h^{-1}$ | Rudge2011 |
| | Random noise in division volume | $\eta_{div}$ | 0.027 | % | Smith2017 |
| | Cell division orientation noise | $\eta_{orient}$ | 0.2 | % | Smith2017 |
| **T6SS attacks** | Attacker firing rate | $k_{fire}$ | 5.0 | $cell^{-1}\ h^{-1}$ | Smith2020 |
| | Dose of effector i translocated per hit | $d_i$ | 0-1 | units | This study |
| | Extracellular needle length | $L_{needle}$ | 0.5 | μm | Smith2020 |
| | Min. needle penetration for hit | $L_{penetration}$ | 0.01 | μm | Smith2020 |
| | Weapon cost per unit secretion | $c$ | 0.0 | h | This study |
| **T6SS response** | Sensitivity to toxin i | $a_i$ | 0.0–1.0 | - | This study |
| | Coefficient of interaction between toxins i and j | $B_{ij}$ | −0.3–0.3 | - | This study |
| | Lysis delay following lethal dose (I=0) | $k_{lysis}$ | 0.125 | h | Smith2020 |
| **Numerical** | Simulation timestep | $\Delta t$ | 0.05 | h | Rudge2011 |
| | Cell / needle sorting grid size | $h$ | 10 | μm | Smith2020 |
| | Conjugate gradient absolute tolerance | $e_{CG}$ | 0.001 | - | Rudge2011 |
| | Max. contact iterations | $Max_{iter}$ | 8 | - | Rudge2011 |
| | Growth restriction factor | $\gamma$ | 500 | - | Smith2017 |

**Table S3: Bacterial strains used in this study**

| Strain | Genotype | Description | Purpose | Source |
|---|---|---|---|---|
| ***A. baylyi*** **ADP1 attackers:** | | | | |
| "*Ab*WT" | *rpsL-K88R, vipA-sfGFP clpV-mCherry2* | Parent ADP1 strain | - | Ringel2017 DH022 |
| "*Ab*Tae1" | *rpsL-K88R vipA-sfGFP clpV-mCherry2 ΔACIAD0053 ΔACIAD1790 ΔACIAD3114 ΔACIAD3425* | ADP1 expressing only effector Tae1 (ACIAD0168) | T6SS Attacker armed with amidase toxin | Ringel2017 PR353 |
| "*Ab*Tle1" | *rpsL-K88R vipA-sfGFP clpV-mCherry2 ΔACIAD0053 ΔACIAD0168 ΔACIAD1790 ΔACIAD3114* | ADP1 expressing only effector Tle1 (ACIAD3425) | T6SS Attacker armed with lipase toxin | Ringel2017 PR305 |
| "*Ab*Tae1Tle1" | *rpsL-K88R vipA-sfGFP clpV-mCherry2 ΔACIAD0053 ΔACIAD1790 ΔACIAD3114* | ADP1 expressing both Tae1 and Tle1 effectors (ACIAD0168, ACIAD3425) | T6SS Attacker armed with both amidase and lipase toxins | Ringel2017 PR317 |
| "*AbΔhcp*" | *rpsL-K88R, Δhcp vipA-sfGFP clpV-mCherry2* | ADP1 with no functional T6SS | T6SS-negative control to test EC adaptation to co-culture | Ringel2017 DH039 |
| ***E. coli*** **MG1655 defenders:** | | | | |
| "AncG" | MG1655::eGFP Kan$^R$ | Kan-resistant ancestral defender strain with eGFP tag | Experimental resistance evolution | Van Der Woude lab (University of York) via Michael J Bottery |
| "AncR" | MG1655::mCherry Kan$^R$ | Kan-resistant ancestral defender strain with mCherry tag | Reference strain for cytometric fitness assay | Van Der Woude lab (University of York) via Michael J Bottery |

### Supplementary Figures

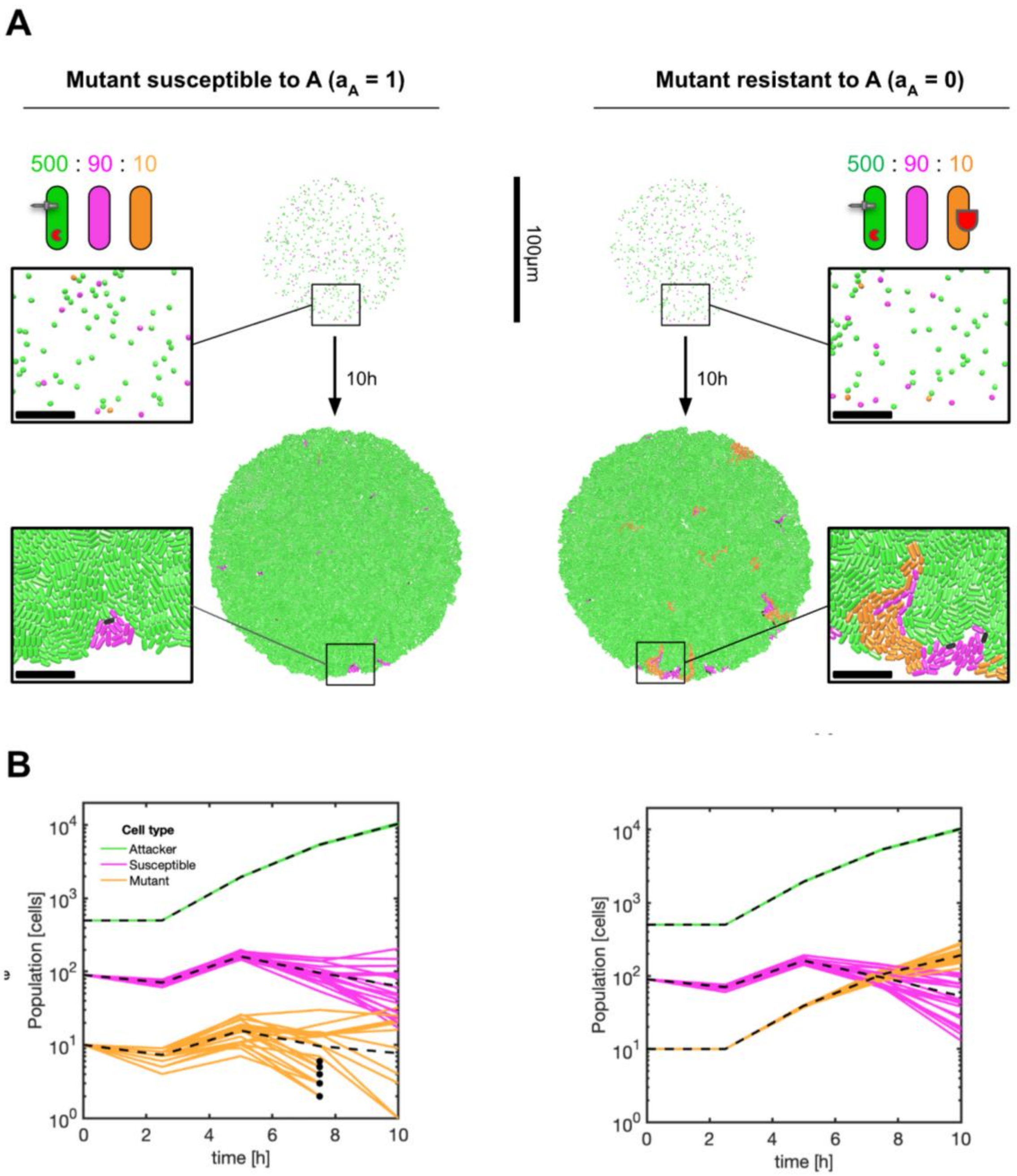


**Fig. S1. Resistance selection simulations using an agent based model.** To quantify evolutionary selection for T6SS resistant-strains, we simulate competitions between T6SS-armed attackers (green), susceptible cells (magenta), and rare mutant cells (orange) with variable sensitivities $a_i$ to attackers' toxins. **(A)** Example competition simulations for mutants having high ($a_A$=1, left column) or low ($a_A$=0, right column) sensitivity to a focal toxin "A" (red pacmen). We initiate simulations by randomly scattering cells in a 500:90:10 (attacker:susceptible:mutant) ratio within a 100μm diameter circle (top row). Cells are then allowed to grow, divide and interact for 10h, before final frequencies of competing cells are assessed (bottom row). Scale bars: 100μm (whole colony images) and 10μm (zoomed sections). **(B)** Example population traces (coloured lines) for the scenario shown in A, indicating absence of selection for a T6SS-sensitive mutant ($a_A$=1, left column), but positive selection for a T6SS-resistance mutant ($a_A$=0, right column). Dashed black lines indicate means of N=20 simulation replicates; black circles mark events where the mutant population goes extinct due to T6SS killing.

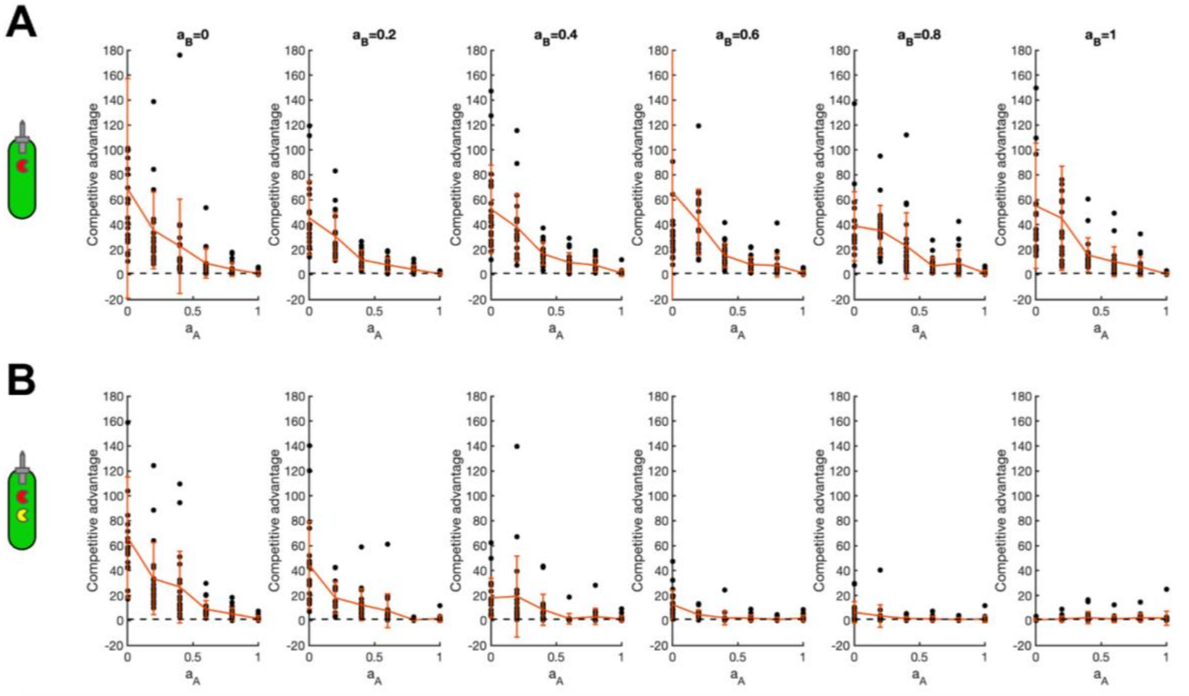


**Figure S2: Multi-toxin attackers limit zone of positive selection for rare T6SS-resistant mutants.** (**A)** Plots of Competitive advantage (without log transformation) chart selection for rare mutants, as a function of resistance to individual toxins ($a_A$, graph axes; $a_B$; columns), against attackers armed only with toxin "A" (red pac-man). Data replotted from Fig. 1D. **(B)** As (A) but against attackers armed with both "A" and "B" toxins (red and yellow pac-man). Data replotted from Fig. 1E. In A, B, black dots correspond to individual simulation replicates; red lines and bars show data means and standard deviations from N=20 0h replicate simulations, each initiated with random assortment of 500:90:10 (attacker:sensitive:resistant) cells. Dashed horizontal lines correspond to absence of selection (Competitive advantage = 1).

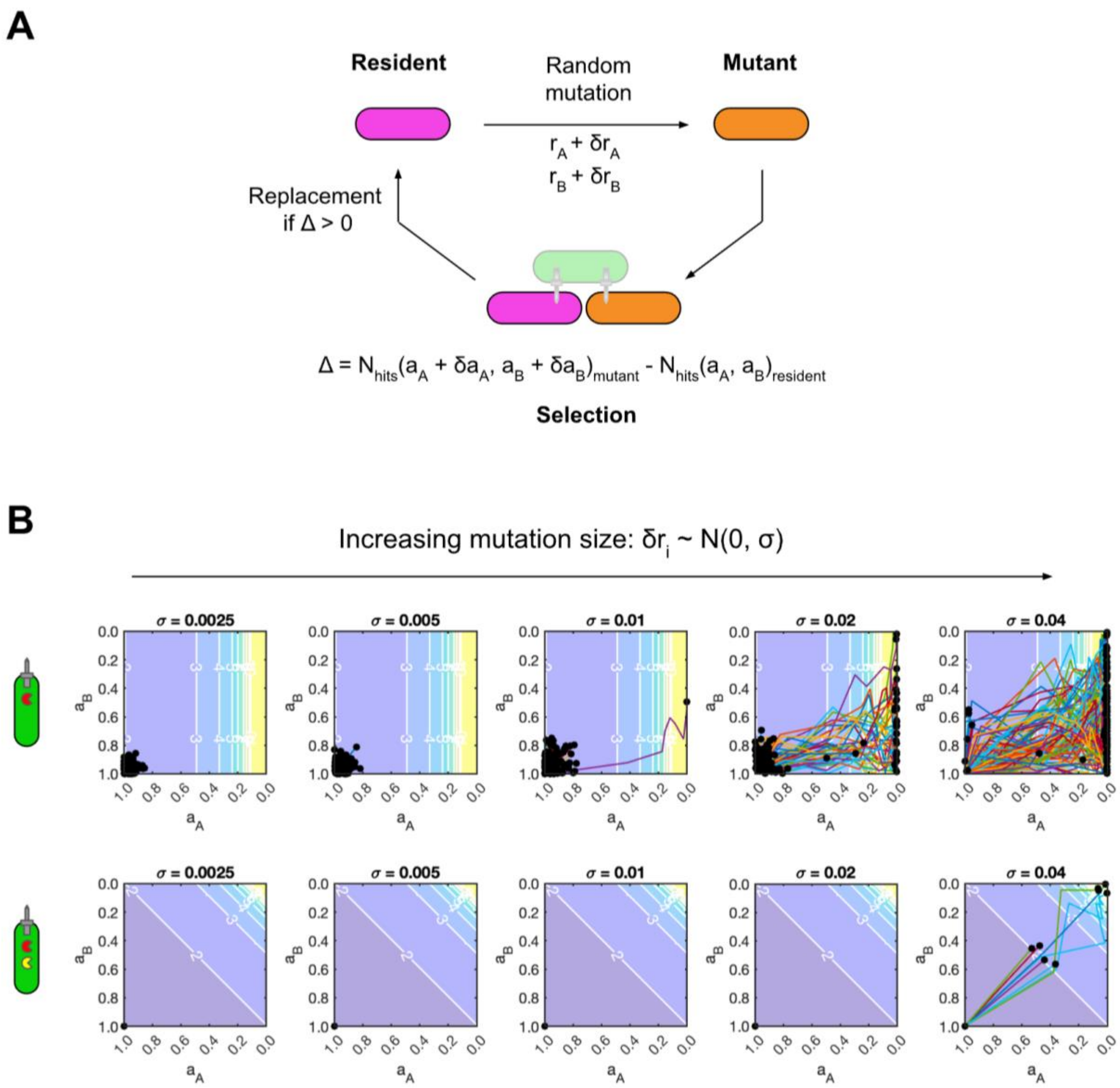


**Fig. S3: Multi-toxin attackers supress resistance evolution at intermediate mutation rates with a simplified trajectory model. (A)** simplified trajectory model replaces (computationally-intensive) agent-based simulations with a simple comparison of phenotypic resistance ($N_{hits}$) for the mutant and resident strain: if, for a given mutation $a_A + \delta a_A$, $a_B + \delta a_B$, $N_{hits}$(mutant) > $N_{hits}$(resident), the mutant replaces the resident. **(B)** Evolutionary trajectories (coloured lines) simulated using the simplified model, imposing selection by single-toxin (top row) or two-toxin attackers (bottom row), for increasing mutation step sizes σ (columns). Trajectories all share the starting point ($a_A$, $a_B$) = (1,1), and are plotted on a contour map showing phenotypic resistance $N_{hits}(a_A, a_B)$ for each coordinate ($a_A$, $a_B$) (see Methods). Trajectory endpoints are shown as black circles. N=100 trajectories per plot, each consisting of 100 sequential mutations.

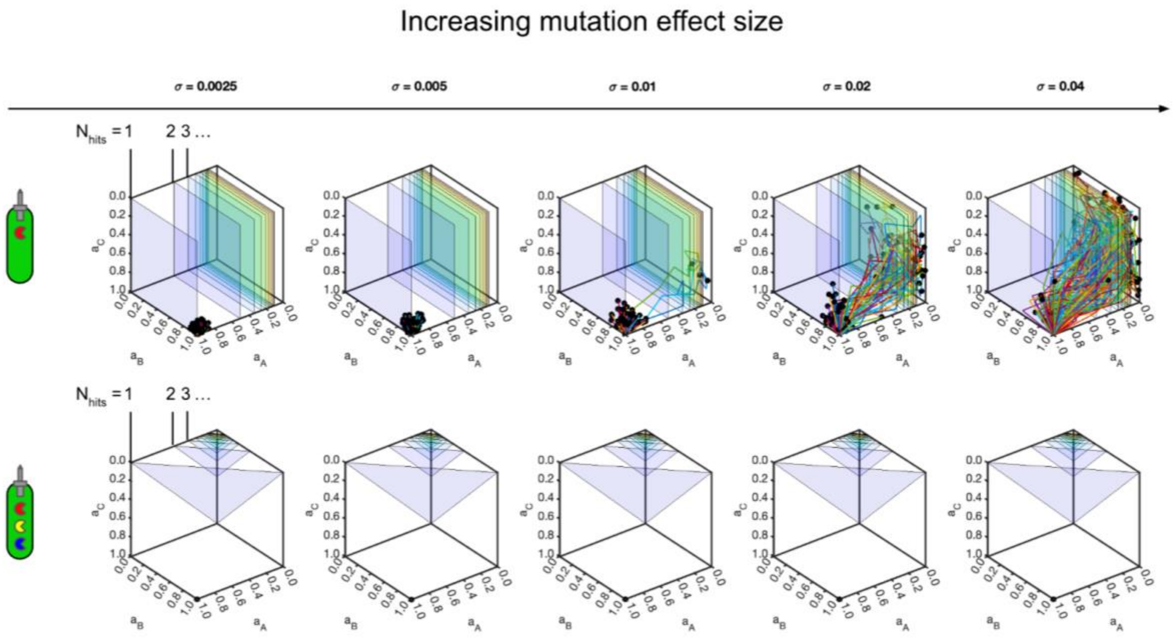


**Fig. S4: Resistance suppression is enhanced by increased toxin arsenal size.** Evolutionary trajectories (coloured lines) for simplified model incorporating three toxins (A, B and C) instead of two, comparing selection by single-toxin (top row) or three-toxin attackers (bottom row), for increasing mutation step sizes σ (columns). Trajectories all share the starting point ($a_A$, $a_B$, $a_C$) = (1,1,1). Coloured transparent surfaces show isoclines in phenotypic resistance ($N_{hits}$ = 1,2,3…10) as annotated. Trajectory endpoints are shown as black circles. N=100 trajectories per plot, each consisting of 100 sequential mutations.

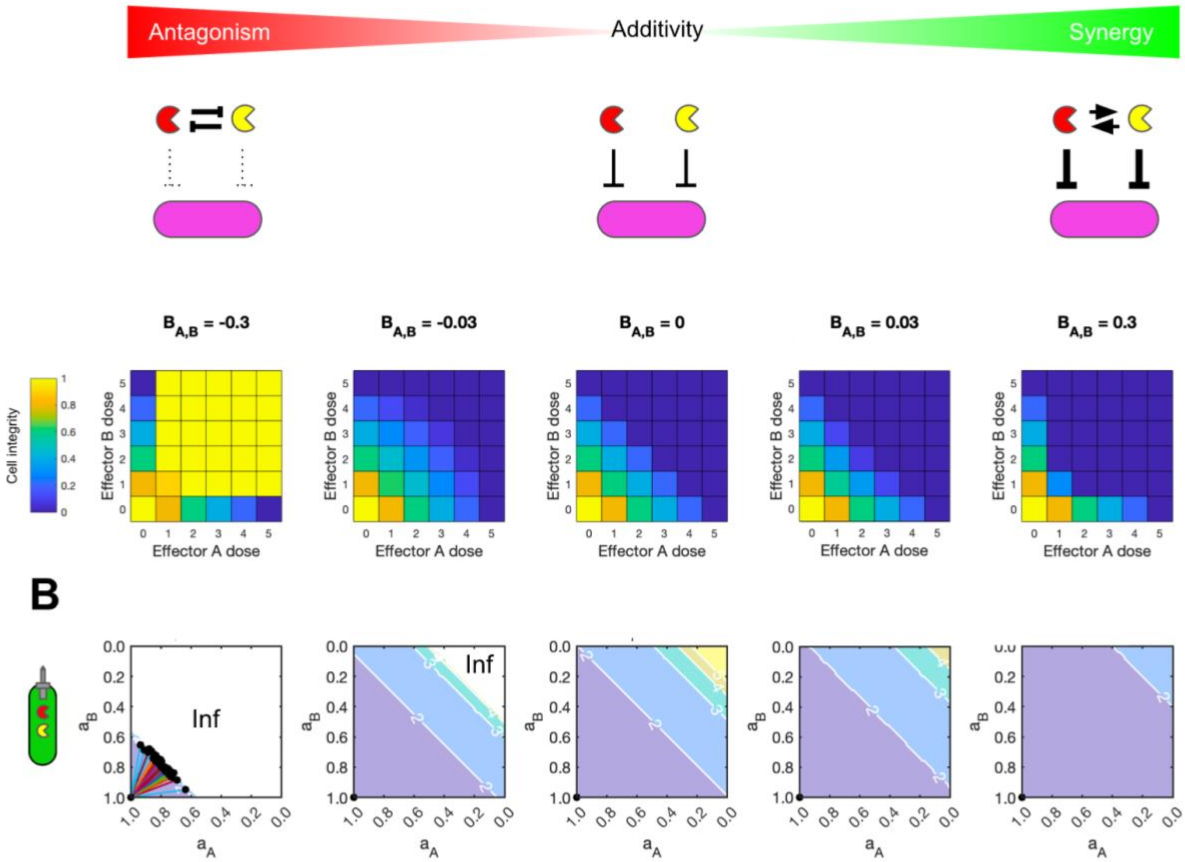


**Fig. S5: Resistance suppression by multi-toxin attackers is enhanced by toxin synergy, but negated by toxin antagonism. (A)** Using our simplified model, we consider the inclusion of antagonistic and synergistic interactions between two T6SS toxins (red and yellow pacmen), examining how these affect overall toxicity and the consequent potential for resistance evolution. Colormaps show residual cell integrity I of a focal cell (with $a_A = a_B = 0.2$) as a function of the dose of each effector, for increasing values of the effector interaction parameter $B_{A,B}$. For $B_{A,B} < 0$ (antagonism, left), the effectors are less toxic together than when secreted individually; for $B_{A,B} > 0$ (synergy, right) the reverse is true. **(B)** Evolutionary trajectories (coloured lines) simulated using the simplified model, imposing selection by two-toxin attackers for the $B_{1,2}$ values shown (columns). Trajectories all share the starting point ($a_A$, $a_B$) = (1,1), and are plotted on a contour map showing phenotypic resistance $N_{hits}(a_A,a_B)$ for each coordinate ($a_A$,$a_B$). Note that in some cases antagonism becomes sufficiently strong that toxicity is completely abolished, such that $N_{hits} = \infty$. Trajectory endpoints are shown as black circles. N=100 trajectories per plot, each consisting of 100 sequential mutations, mutation stepsize $\sigma = 0.1$.

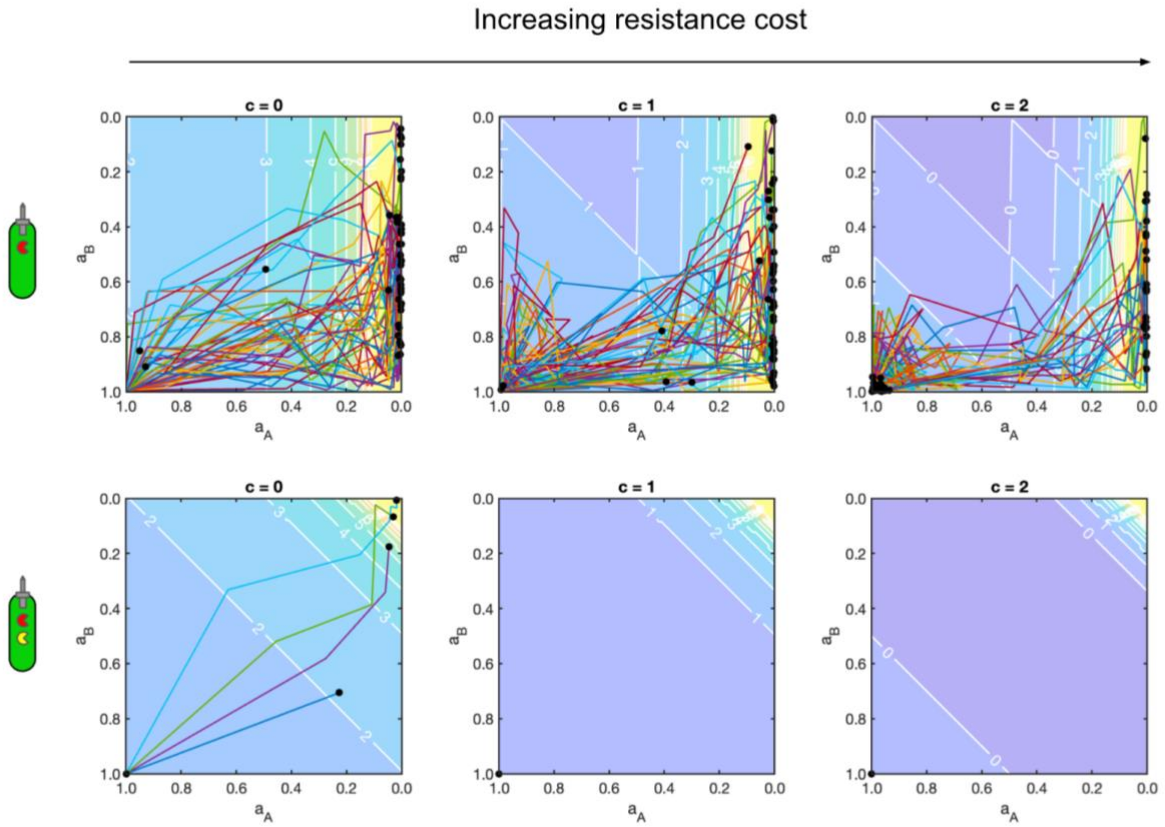


**Fig. S6: Resistance suppression by multi-toxin attackers is enhanced by the addition of resistance fitness costs.** Using our simplified model, we consider the inclusion of fitness costs that scale linearly with resistance to each effector. In this model, a resident or mutant strains' fitness ω is given by $\omega = N_{hits}(a_A, a_B) - c\ (a_A + a_B)$, with parameter c controlling the relative cost of resistance. Above, we show evolutionary trajectories (coloured lines) simulated using the simplified model, imposing selection by single-toxin (top row) or two-toxin attackers (bottom row), for increasing cost values c (columns). Trajectories all share the starting point $(a_A, a_B) = (1,1)$, and are plotted on a contour map showing phenotypic resistance $N_{hits}(a_A, a_B)$ for each coordinate $(a_A, a_B)$. Trajectory endpoints are shown as black circles. N=100 trajectories per plot, each consisting of 100 sequential mutations.

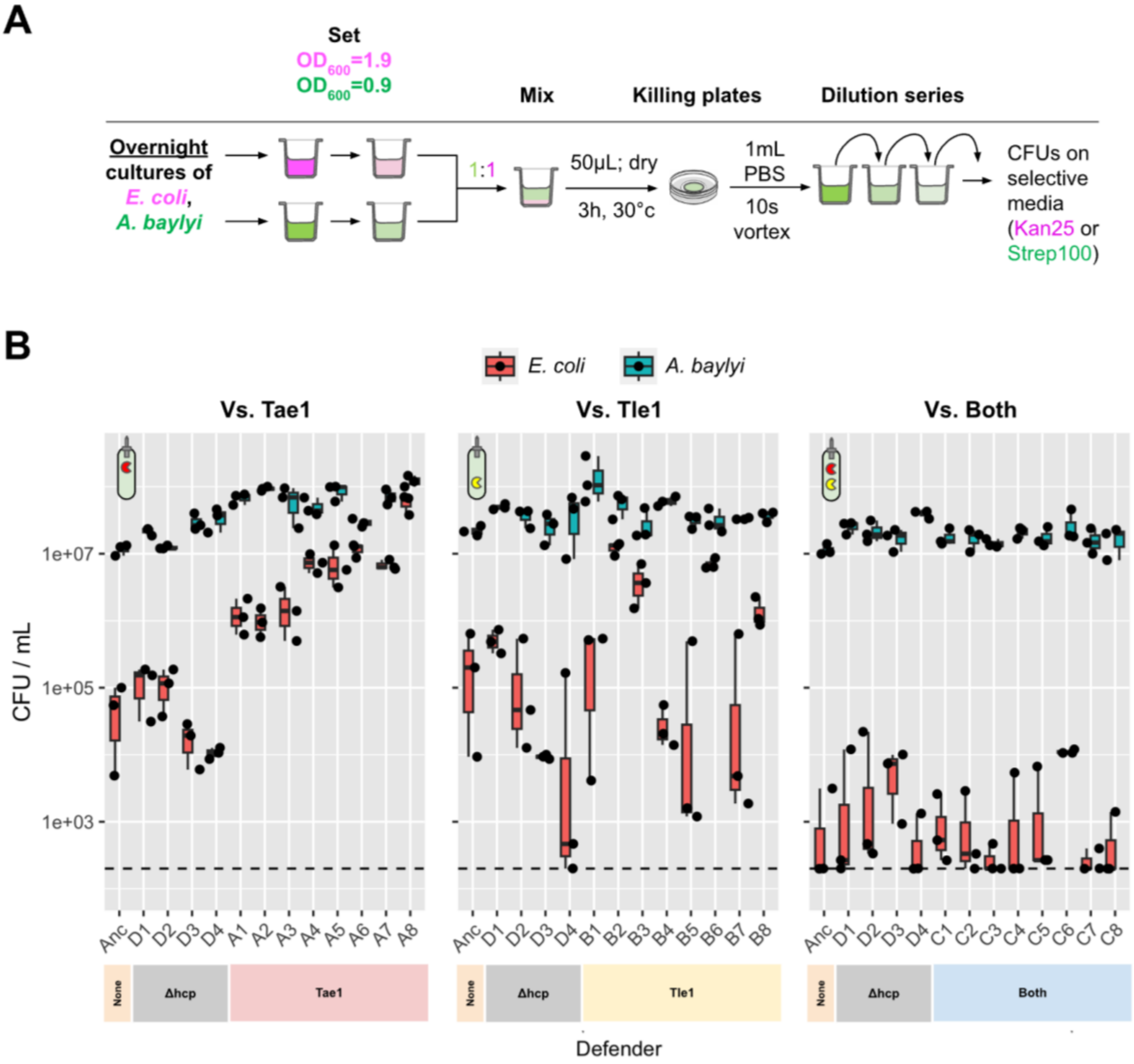


**Fig. S7: Raw recovery data for end-point resistance assay. (A)** Diagram summarising experimental workflow for end-point resistance assay, showing (from left to right) culture density normalisation, mixing, plating, recovery and CFU enumeration. **(B)** Raw CFU data comparing recovery of “Defender” *E. coli* end-point clones D1-4, A1-8, B1-8, C1-8 and the (Anc)estral strain (red), for co-culture with each of the three T6SS attackers (left, middle and right panels). These are data shown alongside *A. baylyi* recovery (teal), plotted as a control for condition uniformity. Box-and-whisker plots show data median, interquartile range and range; individual data points are plotted as black dots. Dashed black line shows the experiments’ detection limit (200 CFU / mL). N=3 pseudobiological replicates; each data point represents the average of 3 technical plating replicates of the same dilution series. The coloured bars below each graph indicate the treatment against which that *E. coli* isolate evolved. *E. coli* recovery data shown are those shown in Figs. 3B (Ancestral strain only) and 3E.

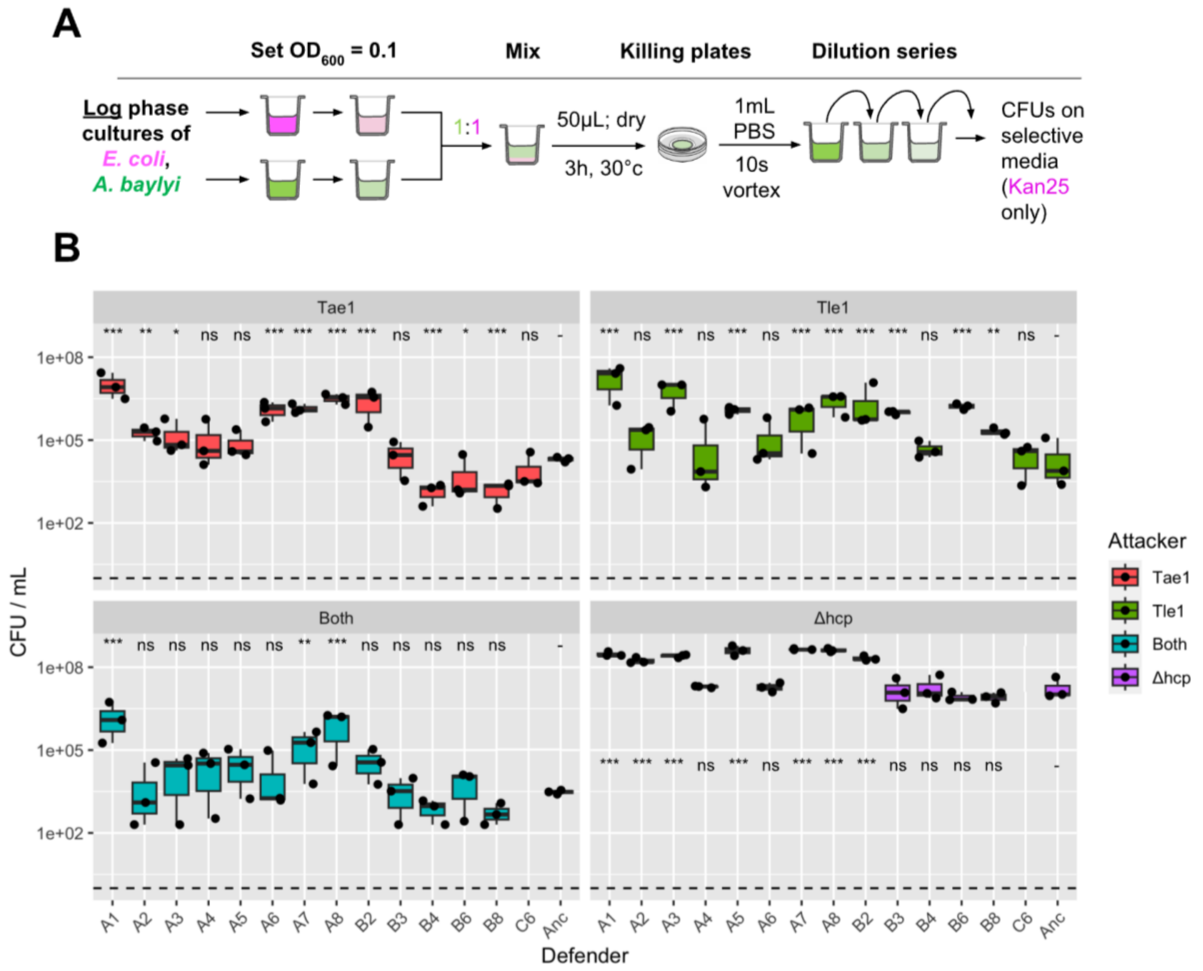


**Fig. S8: Raw recovery data for cross-resistance assay. (A)** Diagram summarising experimental workflow for cross resistance assay. **(B)** Raw CFU data comparing recovery of resistant *E. coli* end-point clones (A1-8, B2-4, B6, B8 and C6 and the (Anc)estral strain, when co-cultured with each of the four T6SS attackers (each shown as a separate panel). Box-and-whisker plots show data median, interquartile range and range; individual data points are plotted as black dots. Dashed black line shows the experiments' detection limit (200 CFU / mL). N=3 pseudobiological replicates; each data point represents the average of 3 technical plating replicates of the same dilution series. Data replotted from Fig. 4A and 4B. Note: isolate C6 was omitted from "Both" and "*Δhcp*" treatments. Significance codes: ***, **, *, ns respectively denote $p<0.001$, $p<0.01$, $p<0.05$ and $p>0.05$; individual isolates assessed for difference with ancestral strain using linear modelling (Dunnett's test for multiple comparisons).

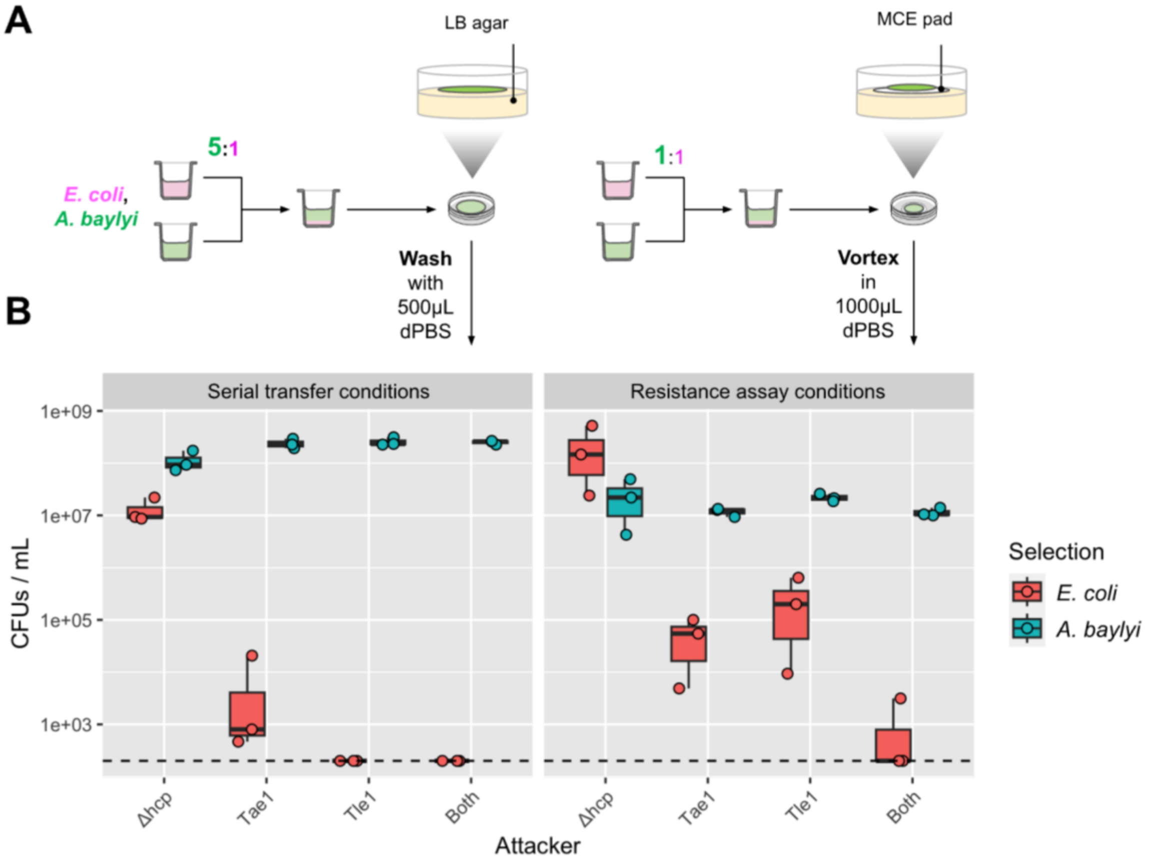


**Fig. S9: Comparison of T6SS toxin lethality between serial transfer conditions and resistance assay conditions. (A)** Protocol comparison between serial transfer experiments (left) and resistance assays (right), which differ in the initial ratio of *A. baylyi* : *E. coli* cells (5:1 or 1:1), the surface on which co-cultures are incubated (LB agar or Mixed [nitro]Cellulose Ester filter pads placed on LB agar), and the method used to recover cells (repeated washing or vortexing). **(B)** Raw CFU data comparing recovery of *E. coli* Ancestral strain (red) and *A. baylyi* attacker strains (teal) as a function of T6SS toxin treatment (x-axes) between these two experimental protocols. Dashed black line shows the experiments' detection limit (200 CFU / mL). N=3 pseudobiological replicates; each data point represents the average of 3 technical plating replicates of the same dilution series. Data shown under "Resistance assay conditions" are replotted from Fig. 3B.
